# Mechanically induced cytoskeletal remodeling in trabecular meshwork cells requires TRPV4 - Rho signaling interactions

**DOI:** 10.1101/2020.08.11.247171

**Authors:** Monika Lakk, David Križaj

**Author notes:** **Corresponding author:** Monika Lakk, 65 N Mario Capecchi Drive, Bldg. 523, Room S4140 JMEC, Salt Lake City, UT 84132. **Author Contributions:** Conception and design of research: M.L. and D.K.; conduction of the experiments: M.L.; data analysis: M.L.; interpretation of results: M.L. and D.K.; preparation of figures: M.L. and D.K; drafting of manuscript: M.L. and D.K., editing and revising the manuscript: M.L. and D.K.; approval of the final version of the manuscript: M.L. and D.K.

## Abstract

Intraocular pressure (IOP) is dynamically regulated by the trabecular meshwork (TM), a mechanosensitive tissue that protects the eye from injury through dynamic regulation of aqueous humor outflow from the anterior chamber of the eye. IOP-dependent increases in TM stiffness and contractility drive open angle glaucoma but the mechanotransduction mechanisms that regulate these processes remain poorly understood. We used fluorescence imaging and biochemical analyses to investigate cytoskeletal and focal adhesion remodeling in human TM cells stimulated with cyclic strain. The cells showed enhanced F-actin polymerization, increased number and size of focal adhesions, and activation of the Rho-associated protein kinase (ROCK). Stretch-induced activation of the small GTPase RhoA, and tyrosine phosphorylations of focal adhesion proteins paxillin, focal adhesion kinase (FAK), vinculin and zyxin were time-dependently inhibited by HC-067047, an antagonist of transient receptor potential vanilloid 4 (TRPV4) channels, and the ROCK inhibitor Y-27632. TRPV4 and ROCK activation were required for zyxin translocation and increase in the number/size of focal adhesions in stretched cells. Y-27632 blocked actin polymerization without affecting calcium influx induced by membrane stretch and the TRPV4 agonist GSK1016790A. These results reveal that mechanical tuning of TM cells requires parallel activation of TRPV4, integrins and ROCK, with chronic stress leading to sustained remodeling of the cytoskeleton and focal complexes.

## Introduction

The ability of eukaryotic cells to change their cytoskeleton in response to the mechanical environment is critical for a wide range of biological and pathological processes (1, 2). In general, application of tensional forces across the lipid bilayer of nonexcitable cells triggers calcium influx via stretch-activated channels (SACs), with Ca^2+^ signaling mediating activation of myosin light chain kinases (MLCKs), inactivation of MLC phosphatases (MLCPs), disinhibition of caldesmon-dependent myosin ATPases and Rho (Ras homology) GTPases, and activity of nonmuscle myosins (3–5). Calcium-mediated phosphorylation of MLC drives formation of contractile actomyosin units and stress fibers, which exert force on integrins, their cytoskeletal anchoring proteins and modulate downstream pathways such as Rho GTPase-dependent modulation of Rho-associated kinase (ROCK) and LIM kinases (6). SACs, the cytoskeleton and contractility shape the structural and functional plasticity of cells exposed to mechanical stress through focal adhesions (FAs), macromolecular assemblies organized around integrin core that bidirectionally transmit tensile forces between the extracellular matrix (ECM) and the cytoskeleton (6–8). Exertion of mechanical forces across FAs increases the accessibility of the cytosolic integrin tail to actin-binding proteins and induces tyrosine phosphorylations of key FA proteins such as paxillin, focal adhesion kinases (FAKs) and vinculin (7, 9, 10) to augment the cells’ resistance to mechanical stress (‘mechanoreciprocity’). Ridley and Hall (1992) showed that active RhoA induces formation of stress fibers and FAs but whether and how SACs interact with the Rho pathway and FAs in the presence of forces is not known.

The biomechanical milieu in the vertebrate eye is dominated by intraocular pressure (IOP), which maintains its shape, regulates development of ocular tissues and modulates cellular signaling. IOP is dynamically controlled by the trabecular meshwork (TM), a layered tissue composed of contractile smooth muscle (SM)-like cells populating ECM beams that are anchored to the endothelial cells lining the canal of Schlemm. Mechanical stretch evokes TM contractions (11), thereby increasing the tissue resistance to aqueous humor drainage to elevate IOP (12, 13). At the cellular level, TM cells compensate for pressure-evoked strains with remodeling of cytoskeletal and ECM pathways that inhibit their ability to drain aqueous humor (14). This process malfunctions in hypertensive glaucoma, as TM cells lose the capacity for homeostatic regulation and adopt the myofibroblast phenotype characterized by increased actin polymerization, α–smooth muscle actin (αSMA) expression, deposition of ECM, and transforming growth factor β (TGFβ) release (14–17). Overall, continued exposure to mechanical stress stiffens the TM and increases its contractility, thereby maintaining IOP in a vicious circle that may result in optic neuropathy (18–20). The remodeling process and IOP elevations in humans and animals with open angle glaucoma can be countered with actin depolymerizers and ROCK inhibitors, which function as IOP-lowering neuroprotective agents in human and animal glaucoma (21, 22). It is therefore of central importance to delineate the molecular mechanisms that drive mechanically induced, Rho-dependent upregulation of TM actin polymerization and contractility.

Here, we tested the hypothesis that force-induced remodeling requires collaboration between transient receptor potential vanilloid isoform 4 (TRPV4), a calcium-permeable polymodal SAC that mediates the responses to strain, swelling, pressure and polyunsaturated fatty acids in TM cells (23–25), and Rho signaling, the actin cytoskeleton and FA pathways. TRPV4 is prominently expressed in mouse and human TM membrane and primary cilia (23, 26) and is often targeted to actin-enriched regions and areas of high strain such as FAs (27–30). Myofibroblast transdifferentiation in smooth muscle, mesenchymal, and endothelial tissues exposed to mechanical stress and/or TGFβ is TRPV4-dependent and bears resemblance to TM remodeling in glaucoma (31–34). In addition, TRPV4 inhibitors protect against fibrosis (33, 34) and lower IOP in a mouse model of glaucoma (23). We found that physiological strain stimuli promote actin polymerization, phosphorylation and/or translocation of key FA-anchoring proteins through a process that requires coordinated activation TRPV4-mediated Ca^2+^ entry and Rho signaling. These findings establish a novel molecular framework that accounts for increased TM stiffness in glaucoma (18), and the efficacy of ROCK/TRPV4 antagonists as IOP-lowering agents in glaucomatous eyes (22, 23).

## Materials and Methods

### Cell culture

Human trabecular meshwork (hTM cells), isolated from the juxtacanalicular and corneoscleral regions of the human eye (ScienCell Research Laboratories, Carlsbad, CA, USA) were cultured in Trabecular Meshwork Cell Medium (ScienCell Research Laboratories, Carlsbad, CA, USA) supplemented with 2% fetal bovine serum (FBS), 100 units/ml penicillin, 100 μg/ml streptomycin, at 37 °C and pH 7.4. In accordance with consensus characterization recommendations (23, 24, 35), the phenotype was periodically validated by profiling for markers *MYOC, TIMP3, AQP1, MGP, ACTA2* (α-smooth muscle actin, αSMA) (Supplementary Figure 6A) and DEX-induced upregulation of myocilin protein (Supplementary Figure 6B & C). The morphology, TRPV4 activation and stretch-induced phosphorylation of FAK and zyxin in hTM cells were virtually indistinguishable from cells dissected from nonglaucomatous donor eyes (56 year old, male and 62 year old, female) (23, 24, 35; Supplementary Figure 7). Human tissues were used in concordance with the tenets of the WMA Declaration of Helsinki and the Department of Health and Human Services Belmont Report. Preliminary accounts of this work were published in abstract form (36, 37).

### Reagents

The TRPV4 agonist GSK1016790A (GSK101) and antagonist HC-067047 (HC-06) were purchased from Sigma (St. Louis, MO, USA). GSK101 (1 mM), Y-27632 (20 mM) and HC-06 (20 mM) DMSO stocks were diluted in extracellular saline, with the final drug concentrations not exceeding 0.1%. The ROCK1/2 inhibitor *Trans*-4-[(1*R*)-1-Aminoethyl]-*N*-4-pyridinylcyclohexanecarboxamide dihydrochloride (Y-27632) was obtained from Cayman Chemical (Ann Arbor, MI, USA) and the calcium indicator Fura-2 AM from Life Technologies (Carlsbad, CA, USA). Salts were purchased from Sigma (St. Louis, MO, USA) or VWR (Radnor, PA, USA).

### Reverse Transcription and Polymerase Chain Reaction (PCR)

Total RNA was isolated using Arcturus PicoPure RNA Isolation Kit according to the manufacturer instructions (Applied Biosystems, Foster City, CA, USA). One microgram of total RNA was used for reverse transcription. First-strand cDNA synthesis and PCR amplification of cDNA were performed using qScript™ XLT cDNA Supermix (Quanta Biosciences. Beverly, MA, USA). The RT-PCR products were run on 2% agarose gels and visualized by ethidium bromide staining, along with 100-bp DNA ladder (Thermo Fisher Scientific, Waltham, MA, USA).

### Quantitative Real-Time Polymerase Chain Reaction (Q-RT-PCR)

SYBRGreen based real-time PCR was performed using Apex qPCR GREEN Master Mix (Genesee Scientific, San Diego, CA, USA). The experiments were performed as triplicates of at least three independent experiments and are expressed as a ~fold change compared to the control. The C_T_ method (ΔΔC_T_) was used to measure relative gene expression where the ~fold enrichment was calculated as: 2 − ^[*Δ*CT (sample) − *Δ*CT (calibrator)]^ after normalization. GAPDH and β-tubulin were utilized as endogenous controls to normalize the fluorescence. The primer sequences are listed in Table 1.

**Table 1.**
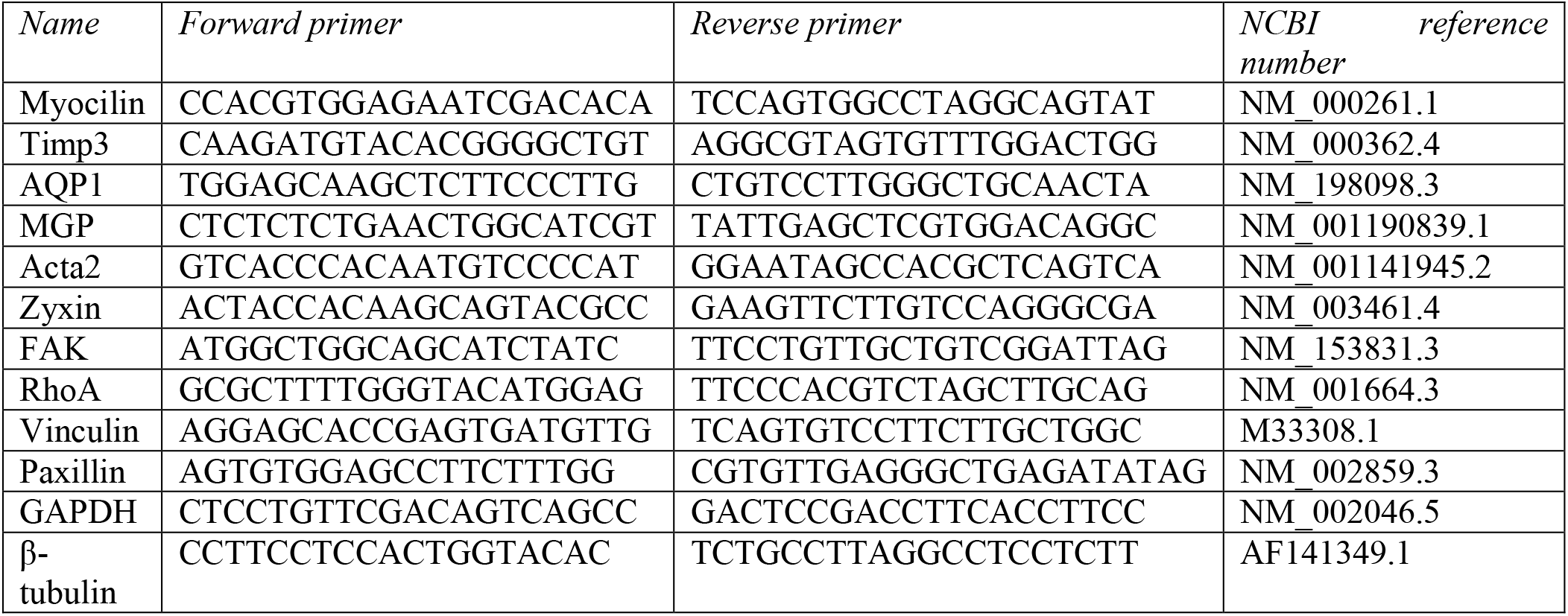
Primer sequences used for PCR analysis.

### Western Blots

Cells were lysed on ice using a radioimmunoprecipitation assay (RIPA) lysis buffer system (Santa Cruz Biotechnology, Dallas, TX, USA) containing PMSF, sodium-orthovanadate and protease inhibitor cocktail (Santa Cruz Biotechnology, Dallas, TX, USA) to avoid degrading processes. An equal amount of protein (20–25 μg) was loaded in 10 or 12 % SDS-polyacrylamide gels. Samples were transferred to polyvinylidene difluoride (PVDF) membranes for 1h at 300 mA. The non-specific binding sites were blocked with 5% non-fat milk and 1% bovine serum albumin (BSA) and incubated overnight at 4 °C with primary antibodies. Primary antibodies and their specifications are listed in Table 2. The binding of the primary antibodies was quantitated with anti-mouse (1:2000, Life Technologies, Carlsbad, CA, USA) or anti-rabbit (1:3000, Cell Signaling, Danvers, MA, USA) horseradish peroxidase (HRP)-conjugated secondary antibodies. HRP conjugated anti-rabbit-GAPDH (1:10.000, Abcam, Cambridge, MA, USA) antibody was used as loading control. The blotted proteins were developed using an enhanced chemiluminescence kit (Thermo Fisher Scientific, Waltham, MA, USA).

**Table 2.**
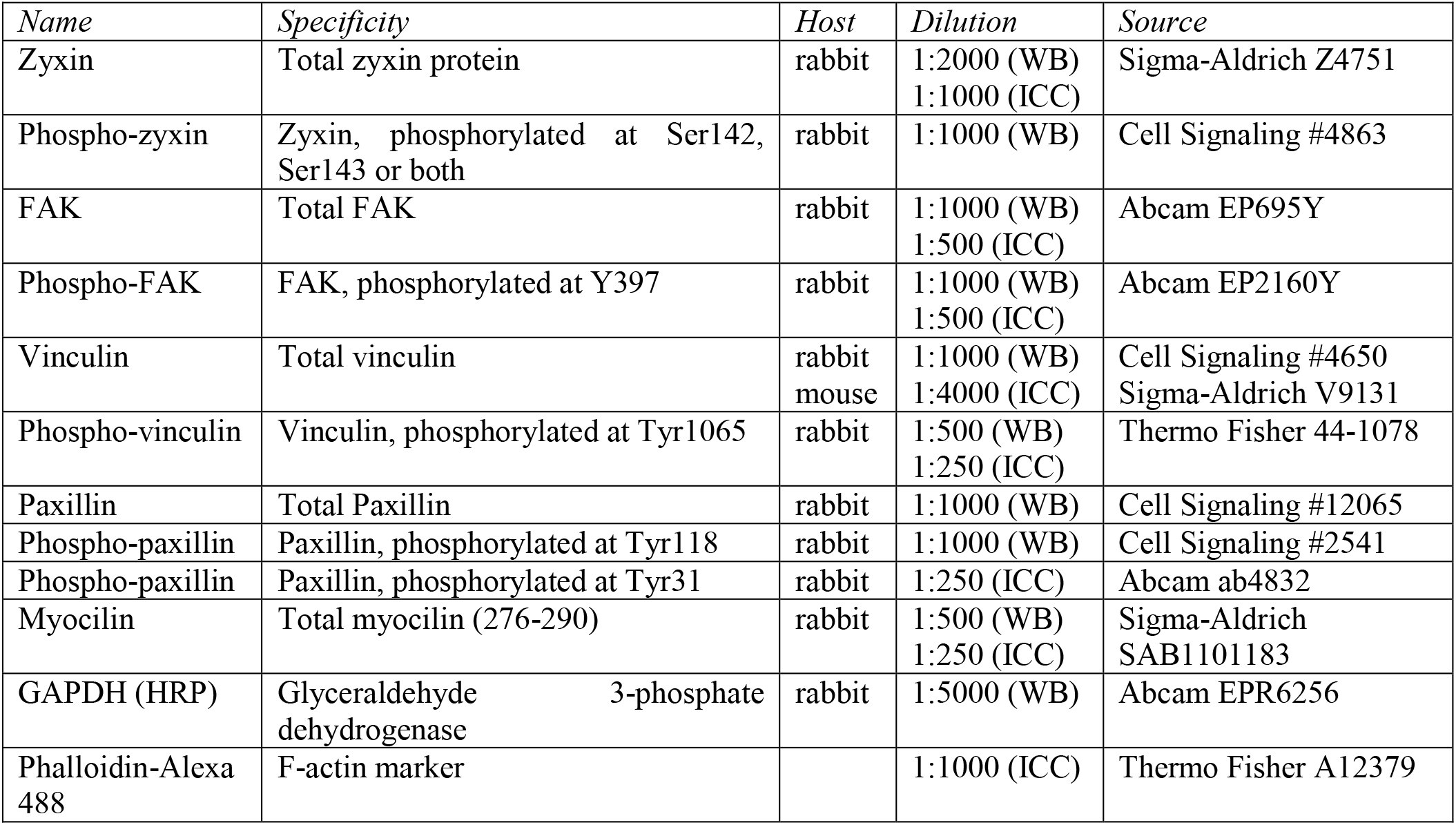
Primary antibodies. *(WB, Western blots; ICC, immunocytochemistry)*

### Rho Activity Assay

GTPase activity was assessed with the Active Rho Detection Kit (Cell Signaling, Danvers, MA, USA) according to the manufacturer’s instructions. Briefly, the assay takes advantage of GTP-bound Rho (active form) binding to the Rhotekin-RBD fusion protein, expressed as a GST-fusion protein in *E. coli*. The protein shows selectivity for GTP-bound as opposed to “inactive” GDP-bound RhoA, RhoB and RhoC isoforms and precipitates active Rho proteins on glutathione resin. The level of activation was quantified with Western blots (the Rho antibody obtained from Cell Signaling was used at 1:500).

### Immunofluorescence

Apoptosis assay: cells were pre-incubated with SRB-VAD-FMK, a cell permeable fluorogenic caspase probe (1:250, AAT Bioquest, Sunnyvale, CA, USA) at 37 °C, 5% CO_2_ for 1 hour before fixation. Cells were fixed with 4% paraformaldehyde for 10 min and blocked in PBS (5% FBS and 0.3% Triton X-100) for 20 min (124, 127). Alexa Fluor 488 phalloidin (Life Technologies, Carlsbad, CA, USA) was utilized for F-actin staining (1:1000 for 1 hr). Primary antibodies (Table 2) were diluted in PBS (2% BSA and 0.2% Triton X-100) and incubated overnight at 4°C. After rinsing, the cells were incubated for 1 hour at RT with goat anti-mouse and goat anti-rabbit IgG (H + L) secondary antibodies conjugated to fluorophores (Alexa Fluor 488 nm, 568 nm and/or 594 nm; 1:500; Life Technologies, Carlsbad, CA, USA). Unbound antibodies were rinsed and conjugated fluorophores protected with Fluoromount-G (Southern Biotech, Birmingham, AL, USA) prior to mounting coverslips. Images were acquired on a confocal microscope (Olympus FX1200) at 1024 × 1024 pixels with a 20X water superobjective (1.00 N.A.; field size: 158.565 × 158.565 μm; 0.155 μm/pixel; sampling speed: 10.0 us/pixel; 12 bits/pixel).

### Immunofluorescence analysis

Images were acquired using identical parameters (HV, gain, offset) resulting very similar signal noise across datasets. ImageJ was used to extract and quantify the mean intensities of immunoreactive signals.

#### Particle counting and colocalization

Particle analysis was performed using ImageJ, with ~40-50 cells per slide averaged across at least 3 independent experiments. Color images were converted to black and white using *Binary*→*Convert to mask* with white background and automatic threshold level. Immunoreactive puncta (number/cell area) with the segmented area were counted with the *Analyze particles* plugin. Minimum (3 pixel^2) and maximum (50-70 pixel^2) pixel area sizes were defined to exclude areas outside of the ROI, calculate the particle number/cell area and determine the relative particulum numbers. Individual particulum sizes for each cell were averaged and normalized. The particulum size was not maximized for p-zyxin analysis in order to capture the particles that translocated along the stress fibers. Colocalization (i.e., overlap of fluorescence for each pixel) was determined for images (Color 1/protein 1 and Color 2/protein 2) that were converted to black/white (*Binary*→*Convert to mask*). To visualize only the colocalized particles, a window was generated with *Process*→*Image calculator* and the *multiply* panel. Colocalized pixels were counted following the subtraction of noise as described above. Minimum pixel size (2 pixel^2) and maximum area (50-70 pixel^2) were determined to exclude objects outside of the area of interest. % colocalization corresponds to the overlap in particulum number/cell area for Colors 1 and 2.

#### Strain application

Strain (substrate deformation) was applied as described (23, 38), using the computer-controlled vacuum-operated Flex 5000 Jr. Tension System (Flexcell International Corporation, Burlington, NC, USA). Cells were plated onto collagen-coated silicon membranes and exposed to cyclic biaxial stretch (6% elongation, 0.5 Hz) for indicated times (1, 3, 5 or 7 hours). Control cells were cultured under the same conditions, but no mechanical force was applied. Y-27632 (10 μM), GSK101 (5 nM) or the HC-06 (1 μM) were added to culture media 30 min prior to stretching. Stretch-induced Ca^2+^ influx was studied in cells loaded with Fura-2-AM (5-10 μM) for 60 minutes and stimulated with biaxial stretch (10% elongation, 1 Hz, 15’ or 30’). The cells were imaged with an upright Nikon E600FN microscope and a 40x (0.8 N.A. water) objective. 340/380 nm excitation was provided by a Xenon lamp within a Lambda DG4 (Sutter Instruments, Novato, CA, USA) wavelength switcher.

#### Calcium imaging

Pharmacological experiments were conducted in fast-flow chambers (Warner Instruments, Hamden, CT, USA) on cells loaded with 5-10 μM Fura-2-AM (K_d_ at RT = 225 nM) for 45 min (23, 128). The saline (pH 7.4) contained (in mM): 98.5 NaCl, 5 KCl, 3 MgCl_2_, 2 CaCl_2_, 10 HEPES, 10 D-glucose, 93 mannitol. TRPV4 and Rho reagents were placed into reservoirs connected to a gravity-fed perfusion system (Warner Instruments, Hamden, CT, USA). Epifluorescence imaging was performed on inverted Nikon microscopes using 40x (1.3 N.A., oil or 0.80 N.A., water) objectives, with excitation for 340 nm and 380 nm filters (Semrock, Lake Forest, IL, USA) delivered by a 150W Xenon arc lamp (DG4, Sutter Instruments) and 2 × 2 binning. Fluorescence emission was high pass-filtered at 510 nm and captured with cooled EMCCD cameras (Photometrics, Tuscon, AZ, USA). ΔR/R (peak F_340_/F_380_ ratio – baseline/baseline) was used to quantify the amplitude of Ca^2+^ signals which were acquired and analyzed using NIS Elements 3.22 software.

#### Plasmid Constructs and Live Actin Imaging

TM cells were transfected with mApple:actin (0.5 μg/ul) or mCherry:actin (0.5 μg/μl) DNA constructs using Lipofectamine (Invitrogen, Carlsbad, CA, USA). Lipofectamine and DNA were diluted separately in Opti-MEM medium; after a quick incubation at RT, the DNA solution was mixed into Lipofectamine solution; the mixture was incubated at RT for about 20 min to obtain the desired N/P ratios. After 24 hour after incubation with the constructs, cells were loaded for 45 min with Fura-2-AM (5-10 μM). TRPV4 and Rho reagents were placed into reservoirs connected to gravity-fed perfusion systems (Warner Instruments, Hamden, CT, USA). *mApple:actin*/*mCherry:actin* and Fura-2 fluorescence were concurrently imaged with a 40x (1.4 N.A. oil) objective (Nikon Ti, Technical Instruments, San Francisco, CA, USA).

#### Statistical Analysis

Data are presented as means ± SEM. For histochemistry, ≥10 per experiment were acquired for at least 3 independent experiments. For calcium imaging, ~10-20 cells per slide were averaged across ~3-6 slides per experiments and 3 independent experiments. Western blot data includes at least 3 separate experiments. Cell numbers = “n”, independent experiments = “N”. The statistical comparisons were determined by one-way ANOVA followed by post-hoc Tukey’s multiple comparison of means (Origin 8.0, Origin Lab Corporation, Northampton, MA, USA). P ≤ 0.05 (*), P ≤ 0.01 (**), P ≤ 0.001 (***) and P ≤ 0.0001 (****) values were considered statistically significant.

## Results

### Stretch-dependent actin stress fiber formation requires TRPV4 activation and Rho signaling

Force-production in non-muscle cells relies on cross-linked actomyosin structures (6, 39). Generally, these are visualized as fluorescently tagged cell-traversing arrays of filamentous actin (F-actin) stress fibers that attach to membrane focal contacts. Physiological levels (6%, 1 hour) of cyclic strain that produce half-maximal [Ca^2+^]_TM_ responses on the stretch-response curve (23) had no effect on shape or viability of TM cells (Supplementary Fig. 1). Indicating that polymerization of TM F-actin is sensitive to mechanical stress, increased tension was associated with significant increases in stress fiber fluorescence (41.7 ± 4.2%; N = 4; P < 0.05) (Fig. 1). The selective TRPV4 blocker HC-067047 (HC-06; 1 μM) did not affect phalloidin signals in resting cells (97.6 ± 4.8%; N = 4) (Fig. 1A & B) but reversed the stress fiber upregulation induced by stretch (to 103.11 ± 4.45%; N = 4; P < 0.01) (Fig. 1A & B). Stress fiber fluorescence in non-stretched cells was not significantly affected by the ROCK inhibitor Y-27632 (10 μM) (88.52 ± 11.65%; N = 4; Fig. 1A & B), whereas ROCK inhibition suppressed the effect of stretch to 71.57 ± 4.81% (P < 0.01; N = 4; Fig. 1A & B). TM cell survival was unaffected by the exposure to HC-06 or Y-27632 (Supplementary Fig. 1B & C). These observations suggest that *de novo* actin polymerization triggered by mechanical stretch requires activation of TRPV4 and Rho signaling pathways.

**Figure 1.**
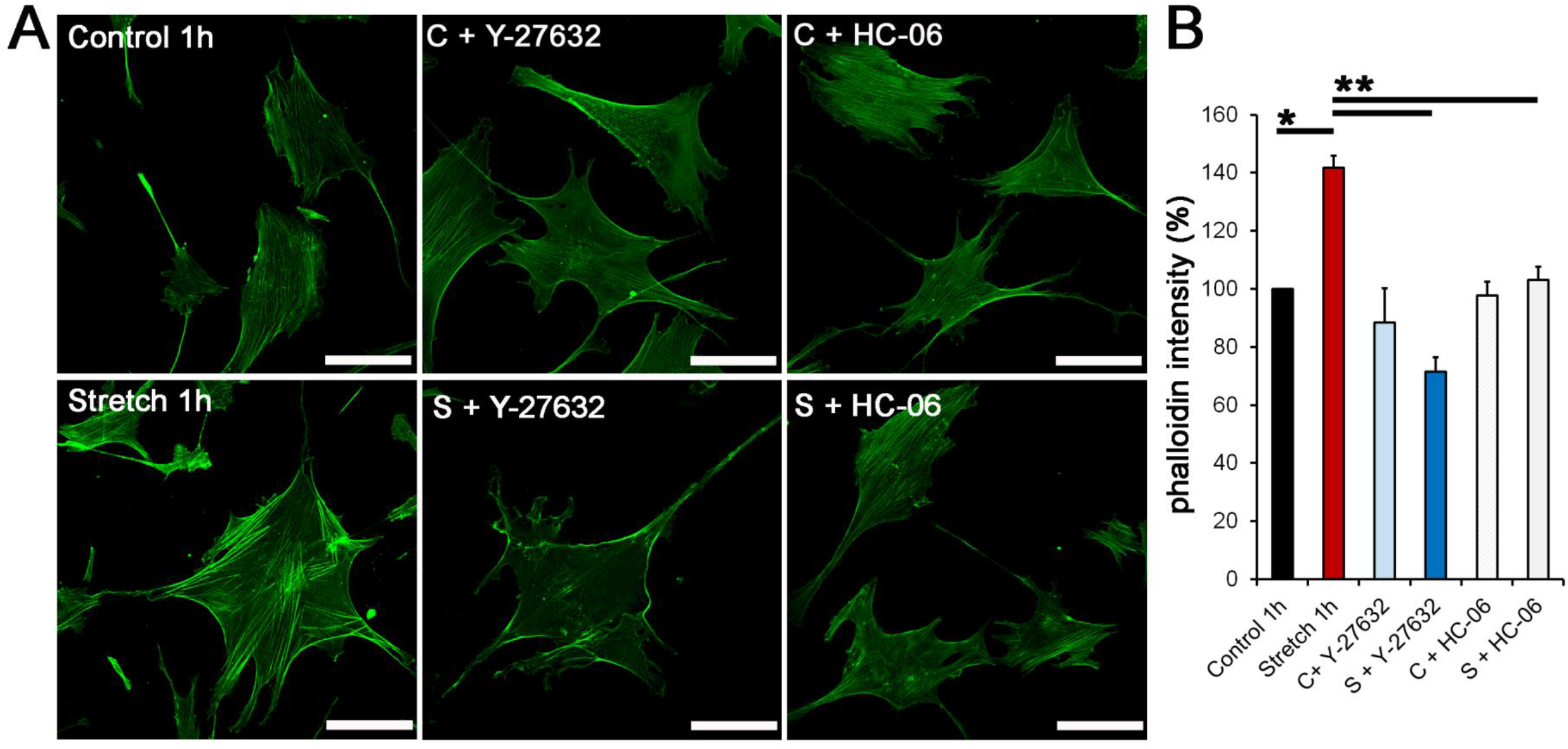
Stretch-induced upregulation of actin cytoskeleton in human trabecular meshwork (hTM) cells requires TRPV4 and ROCK activation. (A) F-actin labeled with phalloidin-Alexa 488 nm and (B) averaged data (N = 4). Preparations were stimulated for 1 hour with 6% cyclic stretch in the presence/absence of TRPV4/ROCK inhibitors. Stretch-induced increases in stress fiber fluorescence were suppressed by HC-06 (1 μM) and Y-27632 (10 μM). Exposure to the antagonists alone had no significant effect on stress fiber formation. Low magnification images are shown in Supplementary Fig. 1A.Scale bar: 20 μm. *, P < 0.05, **, P < 0.01.

### Rho/ROCK signaling pathway is downstream from TRPV4

To gain insight into TRPV4 and ROCK mechanisms that underlie stress fiber formation, we conducted ratiometric calcium imaging in the presence of the TRPV4 agonist GSK101 (5 nM) and/or Rho/TRPV4 inhibitors (Fig. 2A-D). GSK101 elevated ΔR/R to 0.81 ± 0.04 ± 0.27 (n = 164; P < 0.0001; Fig. 2A & D), whereas ROCK inhibition had no significant effect on basal [Ca^2+^]_i_ (0.13 ± 0.01; n = 66; Fig. 2B & D), the amplitude of GSK101-induced calcium signals (0.82 ± 0.06; n = 52; Fig. 2C & D) or F-actin signals (Fig. 1B). GSK101 similarly facilitated the formation of actin stress fibers in cells transfected with *mCherry:actin* or *mApple:actin* (n = 4; Fig. 2Ai-iii). Y-27632 alone non-significantly (Supplementary Fig. 2A) reduced resting stress fiber fluorescence (n = 4; Fig. 2Bi-iii), while concurrent administration of Y-27632 inhibited the activating effect of 5 nM GSK101, resulting in decreased number of stress fibers (n = 4; Fig. 2Ci-iii). In contrast to stretch-induced stress fiber sensitivity to ROCK (Fig. 1), [Ca^2+^]_i_ elevations induced by 15 or 30 min cyclic stretch (to 0.23 ± 0.02 and 0.22 ± 0.02, respectively; n = 30, N = 4; P < 0.0001) were unaffected by Y-27632 (Fig. 2E & Supplementary Fig. 2B), suggesting that Rho-dependent formation of actin stress fibers is downstream from TRPV4. The sensitivity of stretch- and GSK101-induced stress fiber formation to Y-27632 confirms that ROCK is required for actin polymerization driven by both mechanical and chemical inputs.

**Figure 2.**
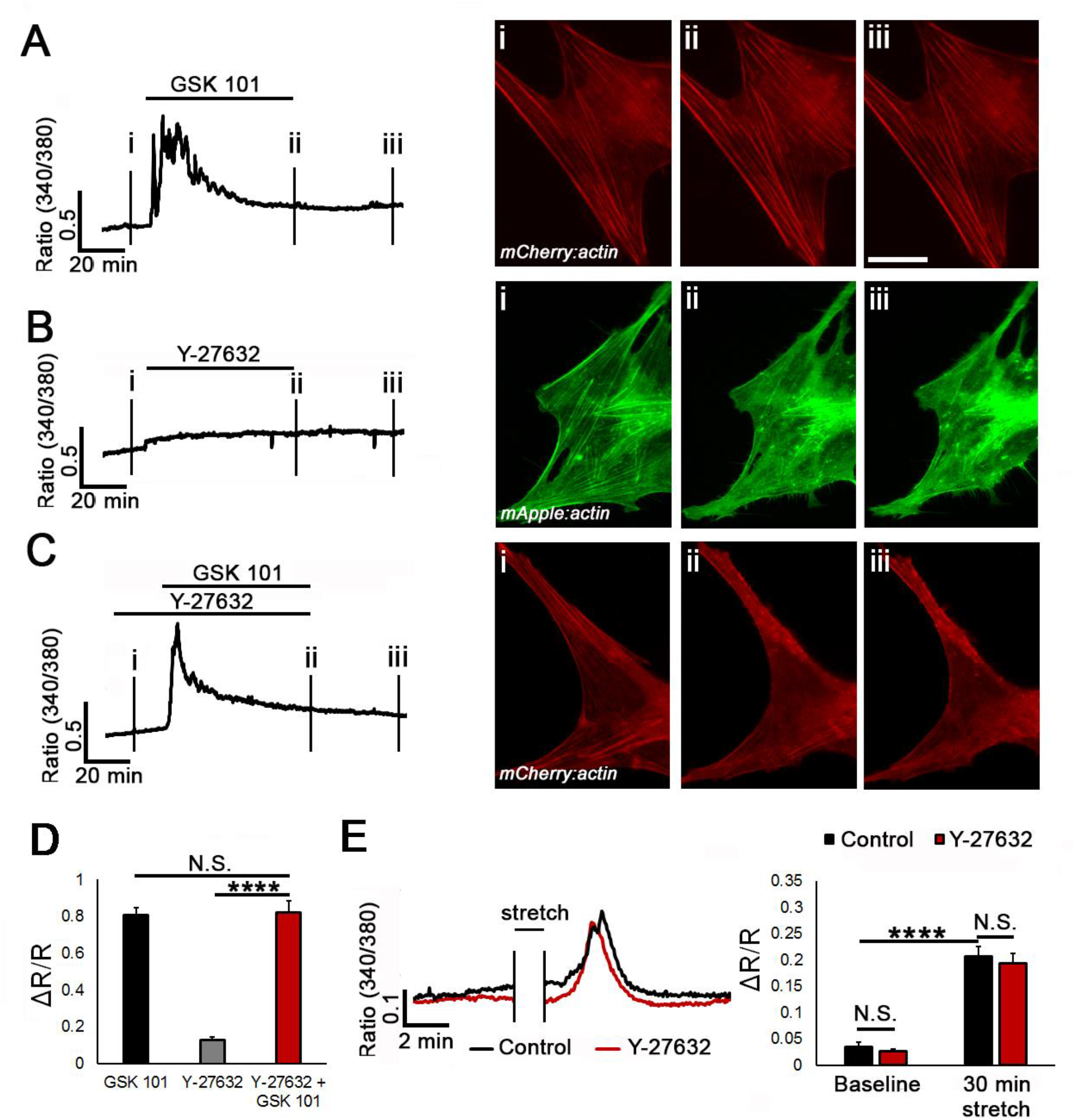
ROCK signaling is downstream from TRPV4. (A - C) Cytosolic Ca^2+^ signals (*left panel*s) and live actin fluorescence (*right panels*) were measured concurrently in Fura-2 AM-loaded cells. (A) GSK101 induced [Ca^2+^]_i_ elevations and reversibly stimulated stress fiber formation (panel ii) in the representative cell (n = 4). (Bi-iii) Y-27632 had no significant effect on baseline [Ca^2+^]_i_ and resting stress fiber formation (n = 4) (Supplementary Fig. 2A). (C) Y-27632 does not affect the amplitude of GSK101-evoked [Ca^2+^]_i_ elevations, however (Ci-iii) Y-27632 significantly (Supplementary Fig. 2A) reduces stress fiber fluorescence (n = 4). (D) Averaged [Ca^2+^]_i_ data from Fura-2 AM-loaded cells (n = 52-162, N = 4). (E) Representative traces and averaged results for 30 min stretched (black trace, bars) and Y-27632-treated (10 μM, red trace, bars) TM cells. [Ca^2+^]_i_ responses induced by stretch are independent of Y-27632. ****, P < 0.0001; N.S., non-significant.

### The small GTPase RhoA is activated by stretch in a TRPV4-dependent manner

We next investigated the stretch-dependence of the small GTPase RhoA, an early responder to tension applied to integrins (40, 41) and a prominent regulator of the TM cytoskeleton (19). One hour of stretch resulted in three-fold increase in RhoA mRNA (3.00 ± 0.72; N = 4; P < 0.05) which was downregulated at longer stretch durations (Fig. 3A). This effect was associated with significant increases in levels of the active (GTP-bound) Rho protein (1.56 ± 0.12; N = 3; P < 0.05) whereas GDP-bound protein levels remained unchanged (Fig. 3B & C). HC-06, but not Y-27632, significantly suppressed stretch-induced upregulation of the active Rho protein (0.99 ± 0.05; P < 0.05; Fig. 3B & C). Hence, tensile strain modulates RhoA transcription and activation in a TRPV4 -dependent manner, but without feedback from ROCK.

**Figure 3.**
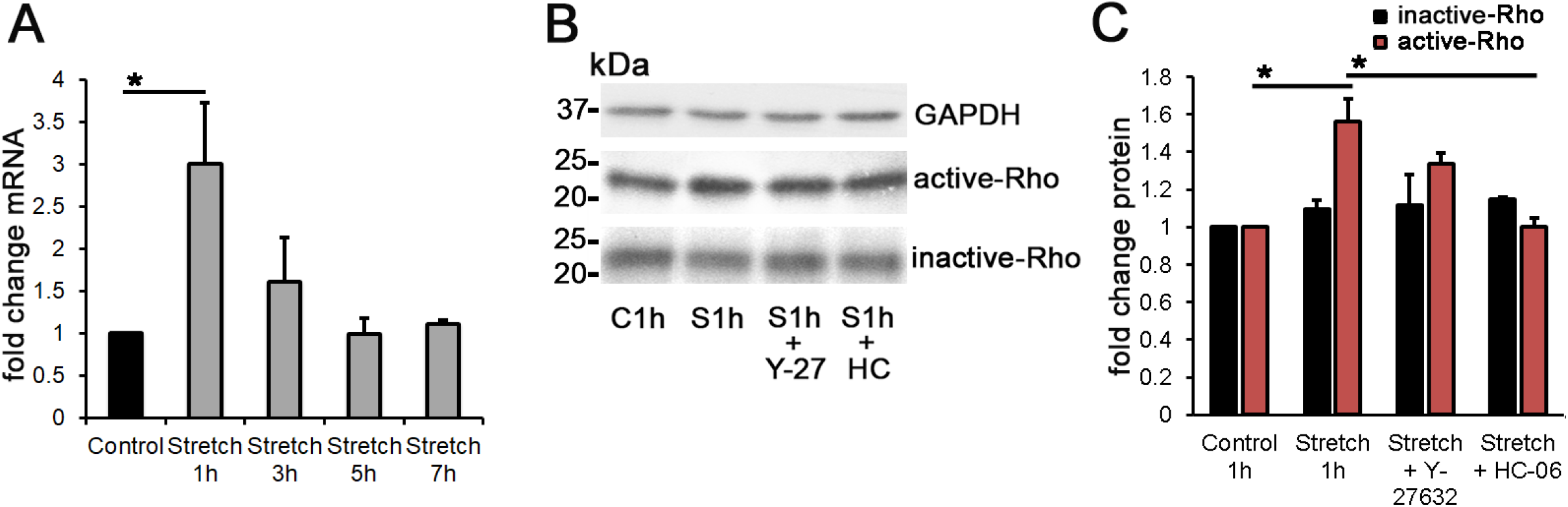
Stretch-induced stimulation of TRPV4 activates RhoA. (A) Semiquantitative RT-PCR, shown as ~fold *RhoA* mRNA following 1 – 7 hours of stretch. *RhoA* mRNA levels transiently increased after 1 hour biaxial stretch (3.00 ± 0.72; N = 4). (B & C) Western blot; active (GTP-bound) Rho protein levels were increased at 1 hour stretch (1.56 ± 0.12; N = 3) whereas the inactive (GDP-bound) Rho levels were unaffected. HC-06 but not Y-27632, suppressed RhoA protein upregulation. *, P < 0.05.

### Stretch-induced phosphorylation and targeting of Focal Adhesion Kinase (FAK) requires time-dependent TRPV4 activation and Rho signaling

Stress fibers generate force by pulling on adhesion sites, which impose stress on the cytoskeleton via adapter and anchor proteins (10, 42). Mechanical strain facilitates formation of FAs, which are enriched in phosphotyrosinated proteins (43). A particularly important anchoring protein, activated by mechanically induced integrin clustering (44, 45), is the Focal Adhesion Kinase (FAK). Phosphorylation of FAK was reported to subserves TM stiffness on rigid substrates (46), regulate phagocytosis (47), myocilin-dependent integrin β1 signal transduction (48) but also maintain the structural stability of actin (49). The Protein Tyrosine Kinase 2 (PTK2) gene that encodes FAK was modestly (1.57 ± 0.12-fold) but significantly (N = 3; P < 0.05) upregulated following 3 hrs strain stimulation and was unaffected at 5 and 7-hour time points (Fig. 4A). We tested whether stretch affects FAK expression and activation by labeling the protein for autophosphorylation at Tyr397 (44, 50). Total FAK protein levels remained constant, but phosphorylation was augmented at 1 and 3 hours to 2.11 ± 0.07; 2.28 ± 0.16 - fold, respectively (N = 3-5; P < 0.01; Fig. 4B). At 1 hour, Y-27632 suppressed phosphorylation (to 0.88 ± 0.14; N = 3; P < 0.05) whereas HC-06 had no effect (at 1.87 ± 0.20; N = 3). At 3-hours stretch, FAK phosphorylation was inhibited by both TRPV4 and ROCK antagonists (Fig. 4C & D). These data suggest that use-dependent formation of FAK-containing focal complexes in the presence of mechanical stress requires time-dependent ROCK and TRPV4 signaling.

**Figure 4.**
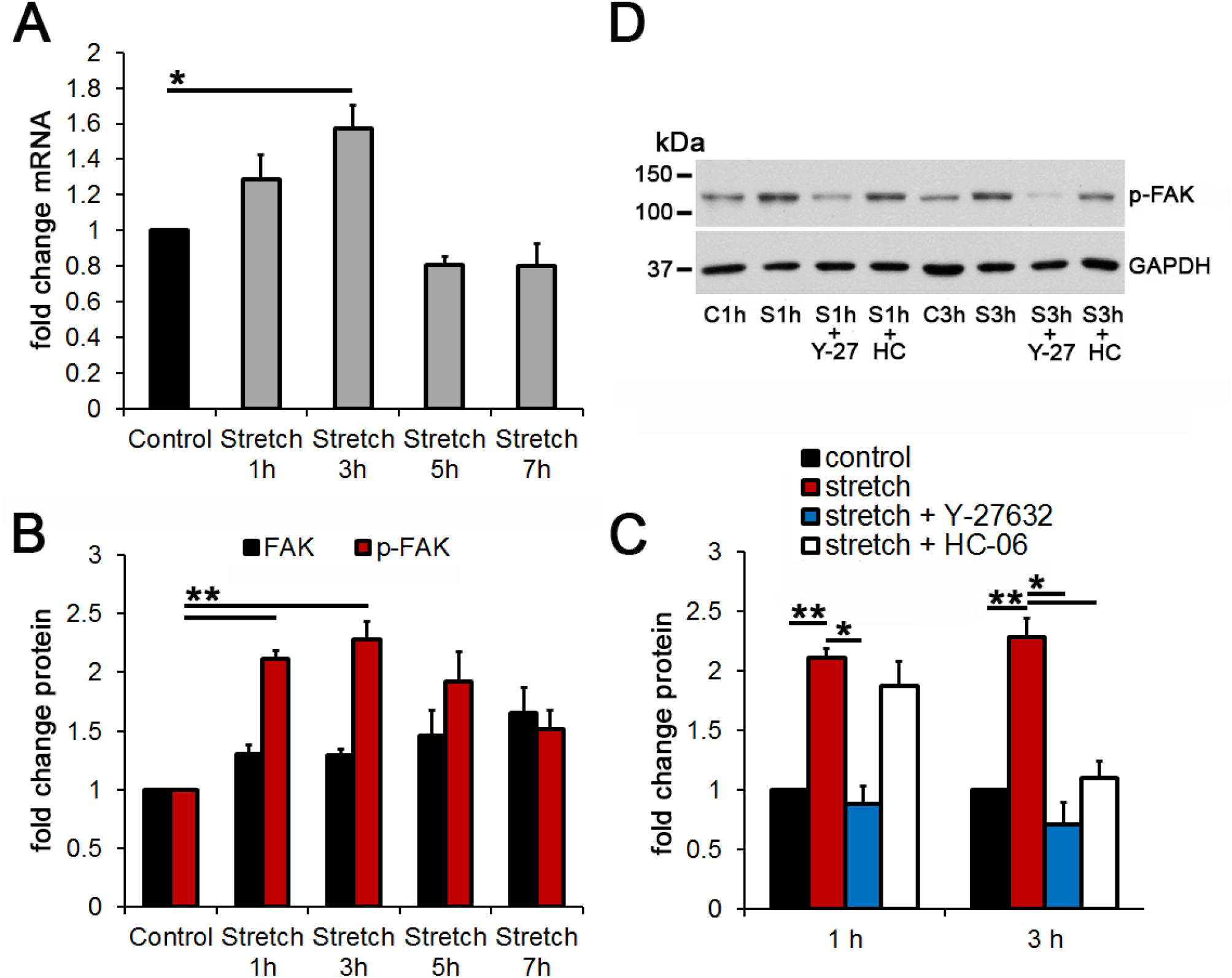
TRPV4 and Rho-signaling time-dependently modulate stretch-induced phosphorylation of Focal Adhesion Kinase (FAK). (A) RT-PCR as ~fold *FAK* mRNA. Strain increases *FAK* mRNA levels at 3 hrs (1.57 ± 0.12-fold; N = 3). (B - D) Western blot data. (B) Total FAK protein levels remain constant, while FAK Tyr397 phosphorylation increases following 1 and 3 hour strain exposure (2.11 ± 0.07 and 2.28 ± 0.16, respectively; N = 3-5). (C & D) pFAK upregulation at 1 hour is inhibited by Y-27632 (0.88 ± 0.14; N = 3) but not HC-06, whereas 3 hour stretch effect is blocked by both antagonists (N = 3-5). *, P < 0.05, **, P < 0.01.

### Transient paxillin phosphorylation is an early step in the TM stretch response

FAK shares the submembrane focal space with paxillin, a 64.5 kDa multidomain scaffolding protein that integrates mechanical cues from the ECM with the integrins (51, 52). Stretch does not affect the levels of paxillin mRNA and total protein at any time point (N = 3; Fig. 5A, black bars in B & C). However, Western blots showed a striking elevation in activated paxillin (Tyr118 phosphorylation) at 1 hour of stretch, with 2.54 ± 0.27 (N = 3; P < 0.01) increase (red bars in Fig. 5B). The effect of stretch was blocked by HC-06 (1 μM; 1.18 ± 0.08; N = 3; P < 0.05) and the calcium chelator BAPTA-AM (100 μM; 0.63 ± 0.06; N = 3; P < 0.01), but not by Y-27632 (10 μM; N = 3; P = 0.216) (Fig. 5Di-ii). These data implicate paxillin activation in early, stretch-and time -dependent TM remodeling that requires TRPV4 and Ca^2+^. Consistent with this, 1 hour stretch caused a remarkable increase in the number of p-paxillin-immunoreactive (ir) (Tyr31-phosphorylated) puncta (to 1.69 ± 0.11; N = 3; P < 0.05 (Fig. 5E & F and Supplementary Fig. 3) without affecting the number of vinculin-ir puncta (Fig. 5F and Supplementary Fig. 3). However, stretch significantly enhanced the extent of vinculin and p-paxillin colocalizations (from 0.20 ± 0.04 to 0.45 ± 0.10 and from 0.51 ± 0.10 to 0.69 ± 0.07, respectively; N = 3; P < 0.05) in a TRPV4-, BAPTA- and ROCK-dependent manner (Fig. 5G and Supplementary Fig. 3).

**Figure 5.**
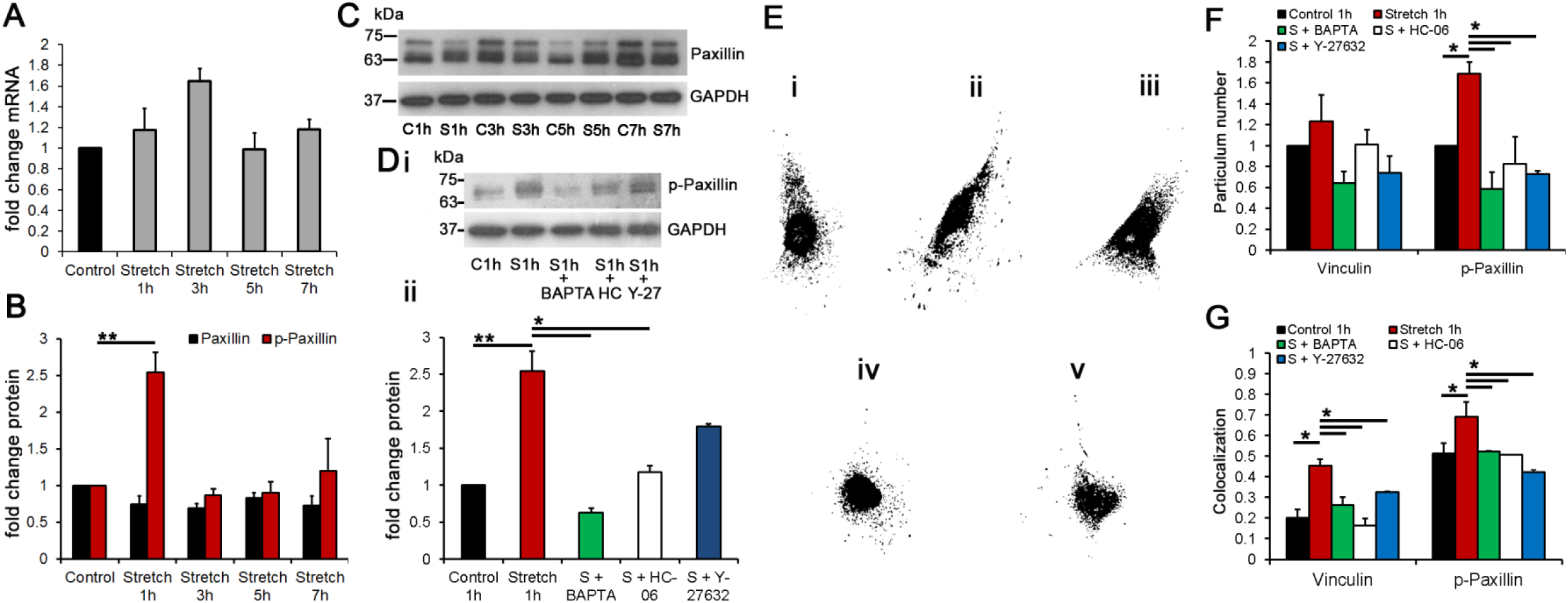
Stretch-induced paxillin phosphorylation requires TRPV4. (A) Semiquantitative RT-PCR, with ~fold changes of *paxillin* mRNA for 1 - 7 hour stretch (N = 3). (B - D) Western blots. (B) Stretch triggers an early elevation of activated (Tyr118) vs. nonactivated forms of the protein (N =3). (C) Western blot for total paxillin in control and stretched samples at 1-7 hours. (Di-ii) Western blot and averaged data for activated paxillin at 1 hour stretch, in the presence of HC-06, Y-27632 or BAPTA. The stretch effect is blocked by HC-06 and the calcium chelator BAPTA (N = 3). (E) Control (i) and stretched (ii) TM cells immunolabeled for p-paxillin, in the presence/absence of BAPTA (iii), HC-06 (iv), or Y-27632 (v). Stretch promotes the dispersal of p-paxillin towards distal membrane sites; TRPV4 and Rho inhibition lowers the extent and dispersion of active paxillin signals. (F) The number of p-paxillin (Tyr31) particuli is upregulated after 1 hour stretch in a TRPV4-, BAPTA- and ROCK-dependent manner, whereas the levels of total vinculin remain unchanged. (G) Colocalization between vinculin and p-paxillin increases in TRPV4- and Rho-dependent manner. *, P < 0.05, **, P < 0.01.

### Stretch promotes colocalization between vinculin and pFAK

Paxillin is recruited to the actin network by the N-terminal domain of vinculin, a key FA protein that regulates integrin and anchor protein dynamics that are required for the enlargement of the focal complex (53). The recruitment of vinculin to stress fiber-anchored FAs was force-dependent (Fig. 6C, F & G). While mRNA and total vinculin protein expression remained unaffected by stretch (Fig. 6A & B), increased colocalization between vinculin and pFAK was observed at 1 and 3 hours (from 46 ± 3% to 58 ± 3% in 1 hr samples and from 57 ± 2% to 71 ± 2% in 3 hr samples; P < 0.05; N = 3) (Fig. 6F & G). This effect was sensitive to Y-27632 at 1 hour (to 39.39 ± 5.04%; P < 0.05; N = 3) and HC-06 at 3 hours (54 ± 4%; P < 0.05; N = 3; Fig. 6F & G). In resting cells, vinculin colocalized with pFAK at the edges of stress fibers (Fig. 6C), with the number of pFAK-ir FA clusters increasing at 1 and 3 hour stretch (by 65.79 ± 20.22% and 34.54 ± 1.01%; P < 0.01 and 0.001, respectively; N = 3; Fig. 6D & E). The increase in immunoreactive puncta was associated with a modest but significant increase in pFAK - vinculin colocalization (from 76.03 ± 2.36% to 85.44 ± 1.25% and from 74.12 ± 1.12 to 86.65 ± 3.1%, respectively; P < 0.05; N = 3) (Fig. 6F & G). Y-27632, but not HC-06, inhibited pFAK upregulation and its interaction with vinculin at 1 hour, whereas HC-06 and Y-27632 both suppressed the effect after 3 hours of stretch (Fig. 6C-G). These findings confirm the Rho-dependence of FAK activation (54) and suggest that TRPV4-dependent regulation of FAK-vinculin-containing FAs involves time-dependent Rho signaling.

**Figure 6.**
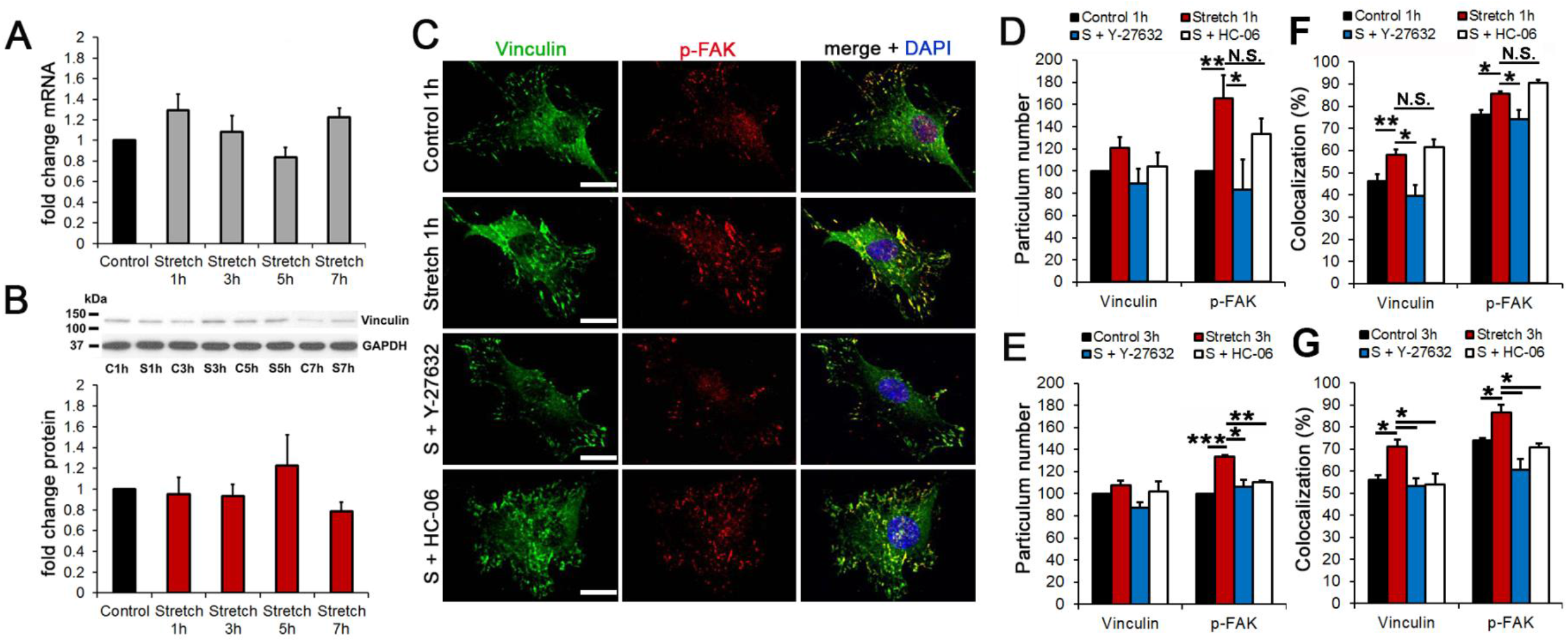
Biaxial stretch facilitates vinculin - pFAK colocalization. (A) Semiquantitative RT-PCR, ~fold changes of *vinculin* mRNA for 1 – 7 hour stretch durations. (B) Western blots. Stretch has no effect on total vinculin protein levels. (C) Control and stretched TM cells in the presence/absence of HC-06 or Y-27632, immunolabeled for total vinculin and pFAK. Averaged data are shown in (D-G). (D & F) 1 hour of stretch upregulates the number of pFAK particuli in a ROCK-dependent manner, and increases the extent of colocalization with vinculin. (E & G) 3 hours of stretch increase pFAK levels and its colocalization with vinculin; this effect is blocked by both HC-06 and Y-27632. *, P < 0.05; **, P < 0.01; ***, P < 0.001. Scale bar: 20 μm.

Stretch also induced increases in vinculin-ir (1.27 ± 0.05; P < 0.01; N = 3) and FA size that were antagonized by ROCK and TRPV4 blockers to 1.01 ± 0.03 and 1.05 ± 0.08, respectively (P < 0.05; N = 3) (Fig. 7A, B and Supplementary Fig. 4A). We tested for stretch-dependence of vinculin activation by labeling for Tyr1065, which is important for force generation (55). A significant increase in activated vinculin (1.78 ± 0.25; P < 0.05; N = 3) was observed following 1 hour of stretch, an effect that was suppressed by Y-27632 (0.75 ± 0.1; P < 0.05; N = 3) and HC-06 (0.99 ± 0.25; P < 0.05; N = 3) (Fig. 7C, D and Supplementary Fig. 4B). Thus, stretch-induced vinculin activation requires Rho and TRPV4 signaling.

**Figure 7.**
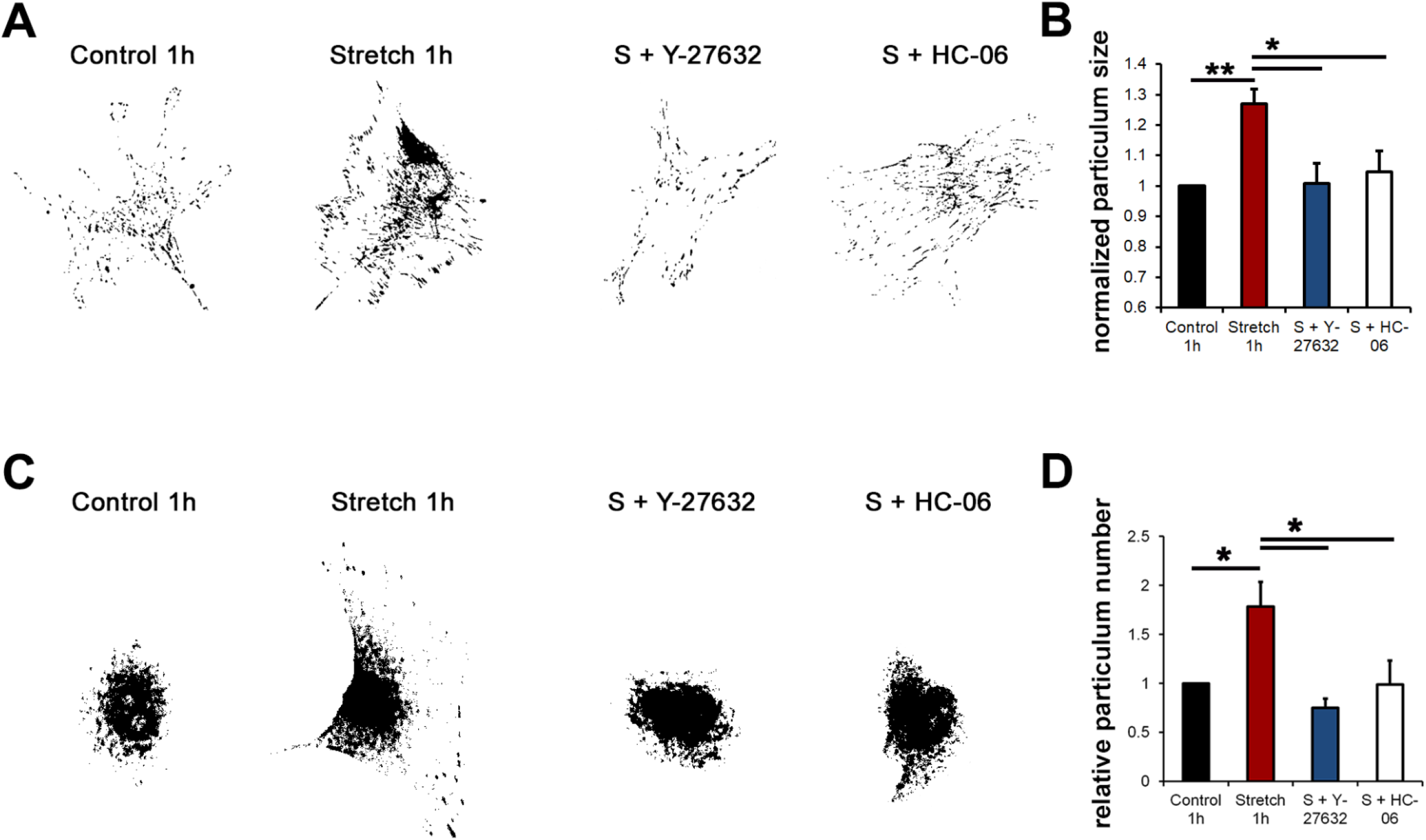
Stretch-induced vinculin phosphorylation requires TRPV4 and Rho signaling. Exposure to strain increases the p-vinculin particulum numbers together with the particulum size of total protein. These effects are suppressed by TRPV4 and ROCK inhibitors. (A) Inverted images of total vinculin immunoreactivity in control, stretch- and antagonist-treated cells. (B) Averaged particulum size of total vinculin in stretched cells in the presence/absence of blockers plotted relative to averaged value in unstimulated cells. The S.E.M. denote variance across separate experiments. (C) Inverted images of Tyr1065-phosphorylated vinculin immunoreactivity in control, stretch- and antagonist-treated cells. (D) Averaged particulum size of activated vinculin in stretched cells in the presence/absence of blockers plotted relative to averaged value in unstimulated cells. N = 3; n = 50-100 cells; *, P < 0.05; **, P < 0.01.

### Stretch-induced translocation of zyxin from FAs to polymerized actin requires TRPV4 and calcium

The FA protein zyxin mediates force-dependent interactions between actin regulatory proteins and actin (56, 57) and is sensitive to TGFβ2, a glaucoma-promoting cytokine known to drive TM actin polymerization (58, 59). Assessment of zyxin mRNA and protein levels after 1 - 7 hours exposure to cyclic strain showed significant upregulation in transcription following 1 hour (2.35 ± 0.19-fold) and 5 hours (2.43 ± 0.22-fold) of stretch (N = 3-4; P ˂ 0.01; Fig. 8A & B), whereas total zyxin protein levels were unaffected for up to 7 hours (2.55 ± 0.16; N = 3; P ˂ 0.05) (Fig. 8B & C). As with other FA adhesome components (Fig. 4 & 5), stretch augmented zyxin phosphorylation (at S142/143) by 2.45 ± 0.48 and 2.69 ± 0.48-fold following 1- and 3-hour stretch, respectively (N = 3-4; P ˂ 0.05 and 0.01; Fig. 8B & C). Zyxin phosphorylation at 1 hour was unaffected by Y-27632, HC-06 or BAPTA (Fig. 8C, D & Ei-ii), whereas TRPV4 and ROCK blockade significantly reduced the levels of phosphorylated-zyxin at 3 hours stretch (1.18 ± 0.31 and 1.32 ± 0.29, respectively; N = 3-4; P ˂ 0.05) (Fig. 8C & D). In unstimulated cells, zyxin-ir was largely confined to FA sites whereas stretch induced a remarkable translocation towards actin stress fibers (Fig. 9 and Supplementary Fig. 5), an effect that was sensitive to HC-06 and Y-27632. Zyxin translocation induced by strain was presumably associated with increased cytoskeletal tension and contractility (60–62).

**Figure 8.**
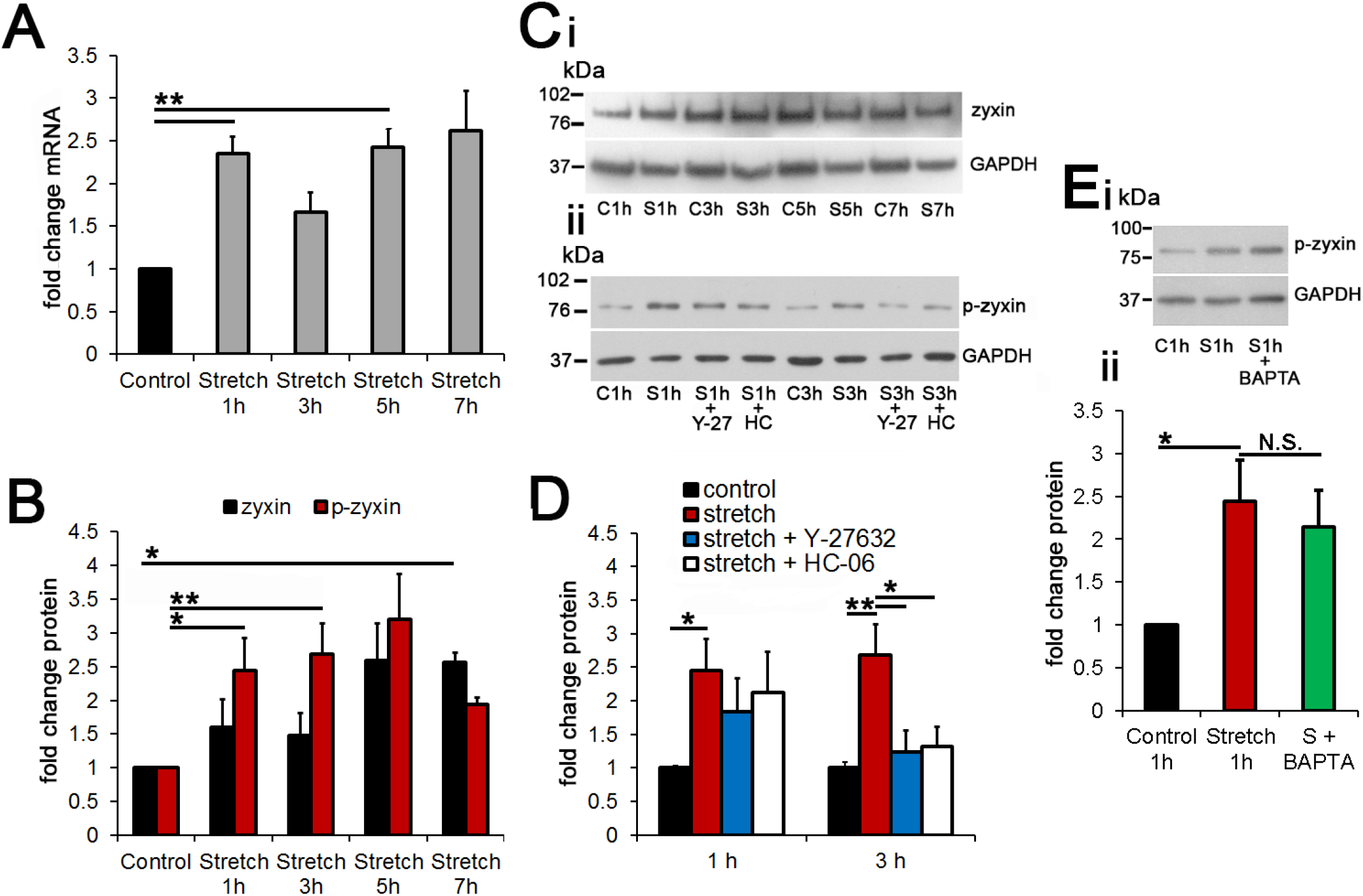
TRPV4 and Rho-signaling time-dependently modulate stretch-induced phosphorylation of zyxin. (A) Semiquantitative RT-PCR. *Zyxin* mRNA is significantly increased at 1 - 5 hours stretch (N = 3-4). (B - E) Western blots. (B) Cyclic stretch triggers increases phosphorylated zyxin protein at 1 and 3 hours (N = 3-4). (Ci) Total zyxin protein in control and stretch samples at 1 – 7 hours. (Cii) Activated zyxin protein in the presence/absence of stretch and TRPV4/ROCK inhibition. Stretch augments zyxin/p-zyxin expression (D) Averaged data for experiments shown in Cii (N = 3-5). Stretch-induced upregulation in p-zyxin is suppressed by HC-06 and Y-27632 at 3 but not 1, hours. (Ei-ii) BAPTA-AM does not affect p-zyxin levels in the presence of stretch (N = 3). *, P < 0.05; **, P < 0.01; N.S. = non-significant.

**Figure 9.**
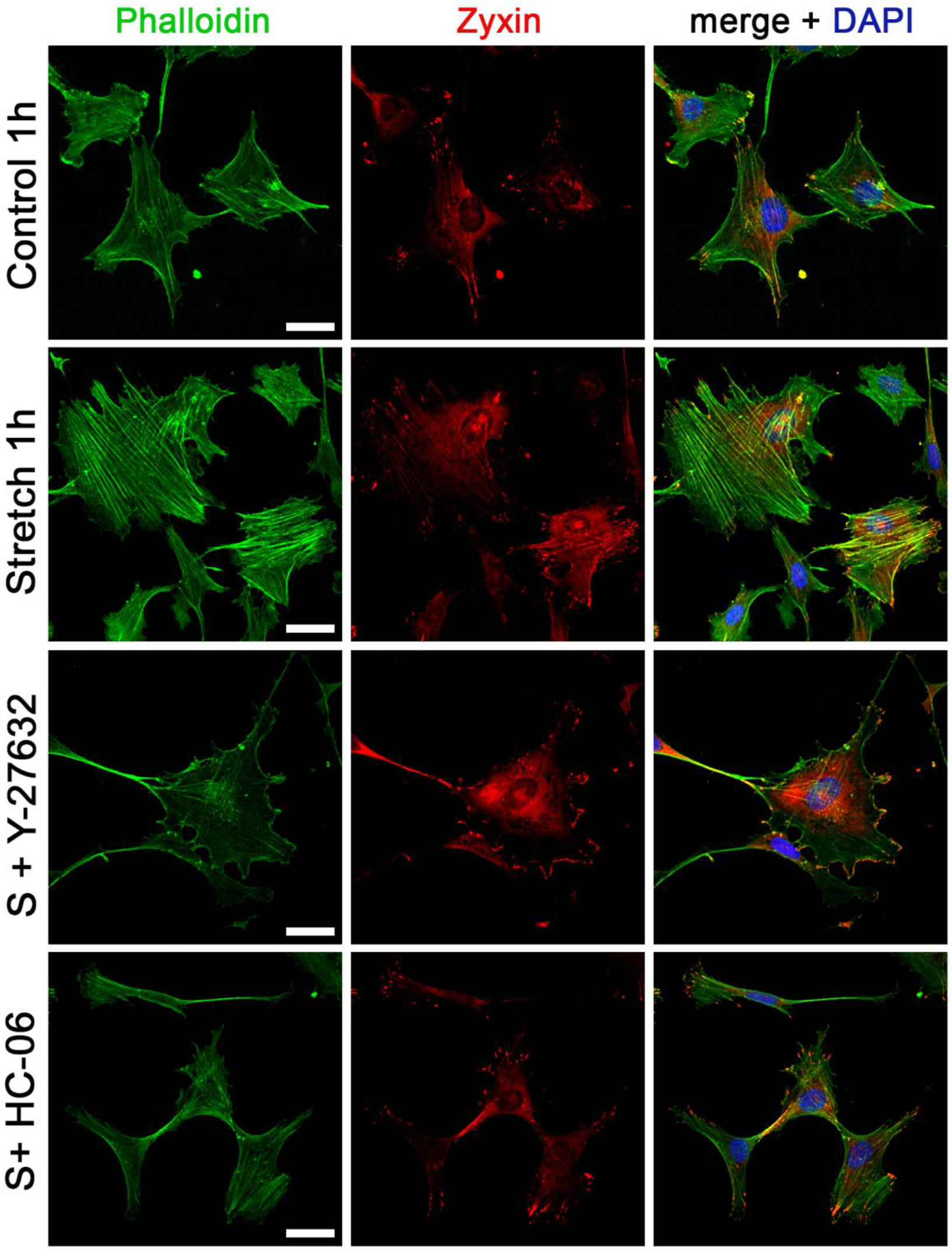
Mechanical force-induced translocation of zyxin requires TRPV4 and Rho signaling. Double labeling for F-actin and total zyxin. 1-hour stretch increases zyxin-ir and promotes its translocation into stress fibers. Stretch-induced translocation of zyxin is inhibited by HC-06 and Y-27632. In the presence of Y-27632, zyxin appears to favors membrane locations (N = 3). Scale bar: 20 μm.

## Discussion

This study identifies the molecular mechanism through which physiological strains regulate the architecture and function of trabecular meshwork cells. We employed strains that mirrored *in situ* biomechanical stress on corneoscleral beams and juxtacanalicular lamellae (~10% –20% strain for pressure differentials of ~5 – 15 mm Hg) (63–65), and a stretch cycle (1 Hz) that reflected ocular pulse dynamics (e.g., 66, 67), to discover that cyclic strains induce cytoskeletal and FA remodeling through obligatory activation of TRPV4 channels, Rho signaling, and TRPV4-Rho interactions. Our data suggest that TRPV4 functions as a likely SAC transducer of TM stretch response and a strong candidate for driving myofibroblast transdifferentiation and cell stiffening associated with stretch- and IOP-dependent increases in F-actin, tyrosine phosphorylation, αSMA & Rho-ROCK signaling and TGFβ release (15, 18, 66, 68–73). Given that strain represents a principal aspect of IOP-generated mechanical force that shapes trabecular outflow (65) whereas cytoskeletal remodeling constitutes the central aspect of glaucomatous TM remodeling (19), elucidation of TM mechanotransduction and Rho signaling pathways brings insight into the biology of the ocular outflow system and identifies pathways that can be targeted to reverse cytoskeletal impairments in glaucoma.

Our results establish a mechanistic framework for how TRPV4 antagonists, actin disruptors and ROCK inhibitors suppress TM stiffness, ECM release and IOP in hypertensive eyes (21–23, 73, 74–77) by adding two important insights: First, Rho signals are downstream from TRPV4, with TRPV4 mechanotransduction; Second, ROCK activation is both necessary and sufficient for increased F-actin polymerization induced by cyclic mechanical strain. By implicating the FA ‘adhesome’ (6) in increased contractility in mechanically stressed TM cells (11, 72), our data provide the molecular context for mechanotransduction-dependent regulation of the outflow pathway. The obliteration of stretch-evoked increases in stress fiber fluorescence by the cell-permeable pyridine derivative Y-27632 (Fig. 1 & 2) is consistent with ROCK functioning as the master regulator of actin polymerization, contractility and remodeling in mechanically stressed TM (22, 74). Rao’s group found that stretch elevates the TM levels of activated RhoA, phosphorylated paxillin and MLCK through aberrant Rho/ROCK activation. Here, we identify TRPV4 as the mechanochannel that is required for dynamic remodeling of F-actin and ROCK-dependent upregulation of integrin-ECM contacts. TRPV4 and ROCK blockers reduced the number and size of activated FAs, and suppressed colocalization between key components of the FA complex (p-paxillin and vinculin, pFAK and vinculin; Fig. 5, 6 & 7), indicating that synergistic contribution of the SAC and ROCK to strain-dependent formation, maturation and maintenance of FAs that reinforces TM cytoarchitecture in the presence of chronic strains.

Membrane stretch promotes F-actin and αSMA expression, MLCK phosphorylation and biosynthesis of ECM that inhibit aqueous outflow (19, 72, 78). Indicating that TRPV4 functions as the SAC that mediates stretch-induced cytoskeletal stiffening, GSK101 mirrored the effect of stretch on stress fiber formation whereas HC-06 blocks stretch-induced F-actin upregulation. ROCK inhibition downregulated pMLCK and αSMA and suppressed F-actin polymerization (66, 72, 75) whereas Y-27632 had no effect on TRPV4-induced [Ca^2+^]_i_ signals (Fig. 2). This indicates that ROCK signaling is downstream from TRPV4-mediated Ca^2+^ influx and rules out a major role for cytosolic or membrane Rho signaling (79) in TRPV4 channel gating, translocation and/or expression. Consistent with this conclusion, the effects of cyclic stretch on cytoskeletal remodeling and myofibroblast transformation can be substituted by either constitutive activation of the Rho pathway (72, 80, 81) or TRPV4 overactivation (31, 34, 82). To the best of our knowledge, our study is among the first to systematically examine the convergence of the two pathways at the level of multiple constituents of the tensile transduction mechanism.

In general, SAC gating is influenced by the properties of the lipid bilayer and by channel interactions with the cytoskeleton and the ECM. The obligatory involvement of TRPV4 in force-induced stress fiber formation, FA reinforcement and fibrosis accords with its functions as a transducer of pressure (24, 83), ECM stiffness (84), cell swelling (38, 85–87), shear stress (88, 89), and its roles in gene expression, integrin signaling (90), tensile activation of vinculin (91) and fibrotic remodeling (34). In contrast to Piezo1 and TREK1 channels which are directly activated by pressure/stretch/indentation (92, 93), the biophysical mechanism that activates TRPV4 channels in the presence of strain is not understood (38, 85, 94–96, 83, 84, 97, 98). Interestingly, β1 integrins activate TRPV4 (5, 30, 90), are expressed in human and mouse TM (99, 100, 101), interact with ROCK (79), and contribute to IOP regulation and glaucoma (102, 103) whereas pretreatment with FAK or β1 integrin inhibitors suppressed sensitization of TRPV4 channels by mechanical stressors (89). Another potential clue is arachidonic acid, which is released by mechanically stimulated TM cells and activates TRPV4 (98, 99). This polyunsaturated ω6 fatty acid might therefore enhance TM sensitivity to weak hydrodynamic loading (23, 96).

Our data identify TRPV4 -mechanotransduction as a venue for Ca^2+^ influx that is necessary and sufficient for stretch-induced reinforcement of stress fibers and growth of focal adhesions. Ca^2+^ activates several FA-associated proteins, including vinculin (91), Src family kinases (SFKs), talin (104) and MLC/Rho-Rho kinases (105). ~30% of [Ca^2+^]_TM_ signal induced by TRPV4 activation is subserved by secondary store release (23), suggesting that the mechanisms studied in the present work may include auxiliary contributions from Ca^2+^ stores and/or store-operated channels (106). Agents that induce Ca^2+^ release from intracellular stores similarly stimulate TM contractility and TGFβ1 signaling to augment the trabecular outflow resistance (35, 80, 107–108).

Our data demonstrate that mechanical stress has dramatic repercussions for the activation state, expression and localization of focal adaptor proteins. This process requires TRPV4 and calcium, as indicated by suppression of stretch-evoked increases in activated RhoA GTPase, paxillin (Tyr118), vinculin (Tyr1065) and zyxin (S142/143) phosphorylation (Figs. 3, 5, 7 & 9) by HC-06 and/or BAPTA-AM. The large early increases in paxillin phosphorylation, its sensitivity to local Ca^2+^ elevations and its localization to FAs point at paxillin as an initial target of increased strain. Consistent with this, TRPV4 inhibition and chelation of calcium suppressed stretch-dependent upregulation in paxillin phosphorylation, number of p-paxillin-ir sites and p-paxillin/pFAK-vinculin colocalization. The increased activation of paxillin and other FA proteins was generally not matched at the transcript level, suggesting that, within the time-frame of our experiments (1 – 7 hours), strain principally regulates FA remodeling through post-translational modifications. Paxillin activation was sensitive to TRPV4-mediated Ca^2+^ influx and ROCK, most likely via Rho-dependent “inside-out” changes in integrin affinity (109). Our finding that FAK migration to vinculin-containing sites and phosphorylation of zyxin acquire TRPV4-dependence after three hours of stretch, suggests that stress fiber enlargement, FA maturation and translocation of zyxin complexes into F-actin stress fibers involve additional Ca^2+^-dependent steps that remain to be investigated. Potential effectors include the Ca^2+^-calmodulin kinase II (CaMKII) which binds integrins upon increases in [Ca^2+^]_i_ (110, 111), prolongs the activation of RhoA (112) and regulates the tensile force in SMCs and myofibroblasts (113), stretch-sensitive Src family tyrosine kinases which bind TRPV4 and integrins, and the protein kinase C, which regulates TGFβ1-dependent upregulation in F-actin and TM contractility (11, 80, 117). A potential caveat on the analysis side is that the substantial increase in the number of immunoreactive puncta in response to stretch might have resulted in the emergence of puncta that were too close for confocal resolution. Thus, we may have underestimated the magnitude of the effect.

A key question in glaucoma research concerns the molecular mechanisms through which elevations in IOP augment the TM resistance to aqueous outflow (65). Previous investigations tended to study the effects of stretch after >8 hours-to-days of stimulation (69, 118–121) however, we found that many key elements of force-induced TM remodeling and contractility are already in place at 1-3 hours of stretch. These include stress fiber upregulation, phosphorylation of RhoA, paxillin, FAK, vinculin and zyxin, and translocation of FA components (zyxin) (Figs. 2–9). Mature FAs contain myosin II directly participate in the contractile response to mechanical stress, which thus involves dynamic cooperation between ECM, membrane and cytoskeletal compartments.

Our findings are consistent with a model whereby TM cells rely on TRPV4 polymodality, Rho GTPases, integrins and Ca^2+^ to integrate parallel aspects of the mechanical and signaling milieu. These mechanisms may be homeostatic at short time scales as IOP fluctuates in response to ocular pulse, eye rubbing, saccades, blinking, swim goggles and yoga poses (65, 67) but become pathological in the presence of sustained stress when TM cells undergo myofibroblast transdifferentiation epitomized by TRPV4-dependent cytoskeletal stiffening, strengthened membrane-ECM contacts and increased contractility. Similar functions were ascribed to TRPV4 in SMC, endothelial, epithelial and mesenchymal remodeling during fibroproliferative cancer, lung and heart diseases (31, 33, 34, 122, 123), suggesting that the channel may participate in a universal program of tissue fibrosis and/or repair. TRPV4-Rho interactions identified here may also be relevant in the hypertensive posterior eye: TRPV4 is expressed in retinal ganglion cells, microglia, Müller cells and endothelial cells (32, 73, 126, 127) and glaucomatous optic nerve head shows elevated Rho GTPase levels (124). In addition to lowering IOP, ROCK inhibitor treatment also improved ganglion cell survival in glaucoma (125). In contrast to Y-2763, which triggers wholesale dissolution of F-actin (Fig. 2), TRPV4 antagonists have no observable side effects (23) and may be used to target the mechanotransducer to maintain function while preserving the cells’ capacity for actomyosin regulation. Because the Rho pathway is downstream of TRPV4, targeting the channel in the anterior and posterior eye could be a viable strategy that combines IOP-lowering with neuroprotection.

## Abbreviations

TM: trabecular meshwork
ROCK: Rho associated protein kinase
FAK: focal adhesion kinase
transient receptor potential vanilloid 4
SACs: stretch activated channels
MLCKs: myosin light chain kinases
MLCPs: MLC phosphatases
IOP: intraocular pressure
ECM: extracellular matrix
αSMA: α-smooth muscle actin
TGFβ: transforming growth factor β
FA: focal adhesion
F-actin: filamentous actin
PTK2: protein tyrosine kinase 2
CICR: Ca^2+^-induced Ca^2+^ release
CaMKII: Ca^2+^-calmodulin kinase II
SMCs: smooth muscle cell
PLA2: phospholipase A2
EET: epoxyeicosatrienoic acid
AA: arachidonic acid
hTM: human trabecular meshwork
FBS: fetal bovine serum
Y-27632: *Trans*-4-[(1*R*)-1-Aminoethyl]-*N*-4-pyridinylcyclohexanecarboxamide dihydrochloride
qRT-PCR: quantitative real-time polymerase chain reaction
RIPA: radioimmunoprecipitation assay
PMSF: phenylmethylsulphonyl fluoride
BSA: bovine serum albumin
HRP: horse-radish peroxidase
ir: immunoreactivity

## Acknowledgements

We thank Dr. Mary Beckerle (Huntsman Center, University of Utah) for actin and zyxin reagents and helpful discussions, and Dr. Amy Lin (Moran Eye Center, University of Utah) and Utah Lions Eye Bank for human tissue.

## Author Contributions

Conception and design of research: M.L. and D.K.; conduction of the experiments: M.L.; data analysis: M.L.; interpretation of results: M.L. and D.K.; preparation of figures: M.L. and D.K; drafting of manuscript: M.L. and D.K., editing and revising the manuscript: M.L. and D.K.; approval of the final version of the manuscript: M.L. and D.K.

## Data Availability

All experimental data are contained within the manuscript.

---

**Supplementary Figure 1. TRPV4 activation and Rho signaling have no effect on TM cell shape or viability.** (A) Representative, low magnification images of cells labeled with phalloidin actin-Alexa 488 nm and DAPI. Y-27632 and HC-06 has no effect on cell and nuclear shape. Stretch upregulates F-actin in a ROCK- and TRPV4-dependent manner without impacting the overall shape (B) Double labeling for F-actin and the fluorogenic SRB-VAD-FMK caspase probe shows rare positive signals seen as cells that lost cytoplasm (white arrow). (C) SRB-VAD-FMK+ signals across cohorts (n = 363-431; N = 2) were normalized to controls. The experimental conditions did not show significant differences from controls. Scale bar: 50 μm.

**Supplementary Figure 2. ROCK signaling is downstream from TRPV4.** (A) Averaged data for experiments shown in Figure 2Ai-iii-Ci-iii. Y-27632 alone had no significant effect resting stress fiber fluorescence, however Y-27632 significantly prevent GSK 101 effect on stress fiber formation (B) Averaged data for 15 min stretched (black bars) and Y-27632-treated (10 μM, red bars) TM cells (n = 30; N = 4). Stretch-evoked [Ca^2+^]_i_ responses are independent of Y-27632. *, P < 0.05; ****, P < 0.0001; N.S., non-significant.

**Supplementary Figure 3. Stretch-induced paxillin phosphorylation requires TRPV4.** Control and stretched TM cells in the presence/absence of BAPTA, HC-06 or Y-27632, immunolabeled for p-paxillin (Tyr31) and total vinculin. The number of p-paxillin (Tyr31) particuli is upregulated after 1 hour stretch in a TRPV4-, BAPTA- and ROCK-dependent manner, whereas the levels of total vinculin remain unchanged. Colocalization between vinculin and p-paxillin increases in TRPV4- and Rho-dependent manner. Averaged data shown in Figure 5F and G. N = 3; Scale bar: 20 μm.

**Supplementary Figure 4. Stretch-induced vinculin phosphorylation requires TRPV4 and Rho signaling.** (A) Immunolabeling for total vinculin protein and (B) for Tyr1065-phosphorylated vinculin. Exposure to strain increases the p-vinculin particulum numbers at 1 hour together with the particulum size of total protein. These effects are suppressed by TRPV4 and ROCK inhibitors. Averaged, normalized data shown in Figure 7B and D. N = 3; *, P < 0.05; **, P < 0.01. Scale bar: 20 μm.

**Supplementary Figure 5. Mechanical force-induced translocation of zyxin requires TRPV4 and Rho signaling.** Averaged data of total zyxin immunofluorescence labeling in Figure 9. One hour stretch increases particulum sizes of zyxin, which suggesting its translocation into stress fibers. This stretch-induced translocation of zyxin is inhibited by HC-06 and Y-27632. (N = 3); *, P < 0.05.

**Supplementary Figure 6. TM cells express standard phenotype and focal adhesion markers.** PCR. (A) Transcripts encoding MYOC, TIMP3, AQP1, MGP and ACTA2 are prominently expressed, together with RhoA, FAK, zyxin, vinculin and paxillin mRNAs; *Gapdh* and ◻*-tubulin,* loading controls. (B) Western blot with an anti-myocilin antibody. 2 day exposure of hTM cultures to 100 nM DEX upregulated myocilin expression (N = 2). (C) Immunolabeling for anti-myocilin antibody after 2 day exposure to 100 nM DEX treatment, showing positive signals in TM cells (N = 2). Scale bar: 50 μm.

**Supplementary Figure 7. TRPV4 and Rho-signaling modulate stretch-induced cytoskeletal reorganization in primary cells.** Western blots of representative marker and cytoskeletal proteins in primary TM cells. (A) Raw data. (B - E) Averaged results (N = 2). Stretch-evoked expression and the effects of HC-06 and Y-27632-in primary TM cells correspond to the effects seen in immortalized cells. Myocilin expression, however, is unaffected by stretch.

