## Supplementary figures and images for "Mechanically induced cytoskeletal remodeling in trabecular meshwork cells requires TRPV4 - Rho signaling interactions"

### Supplement info

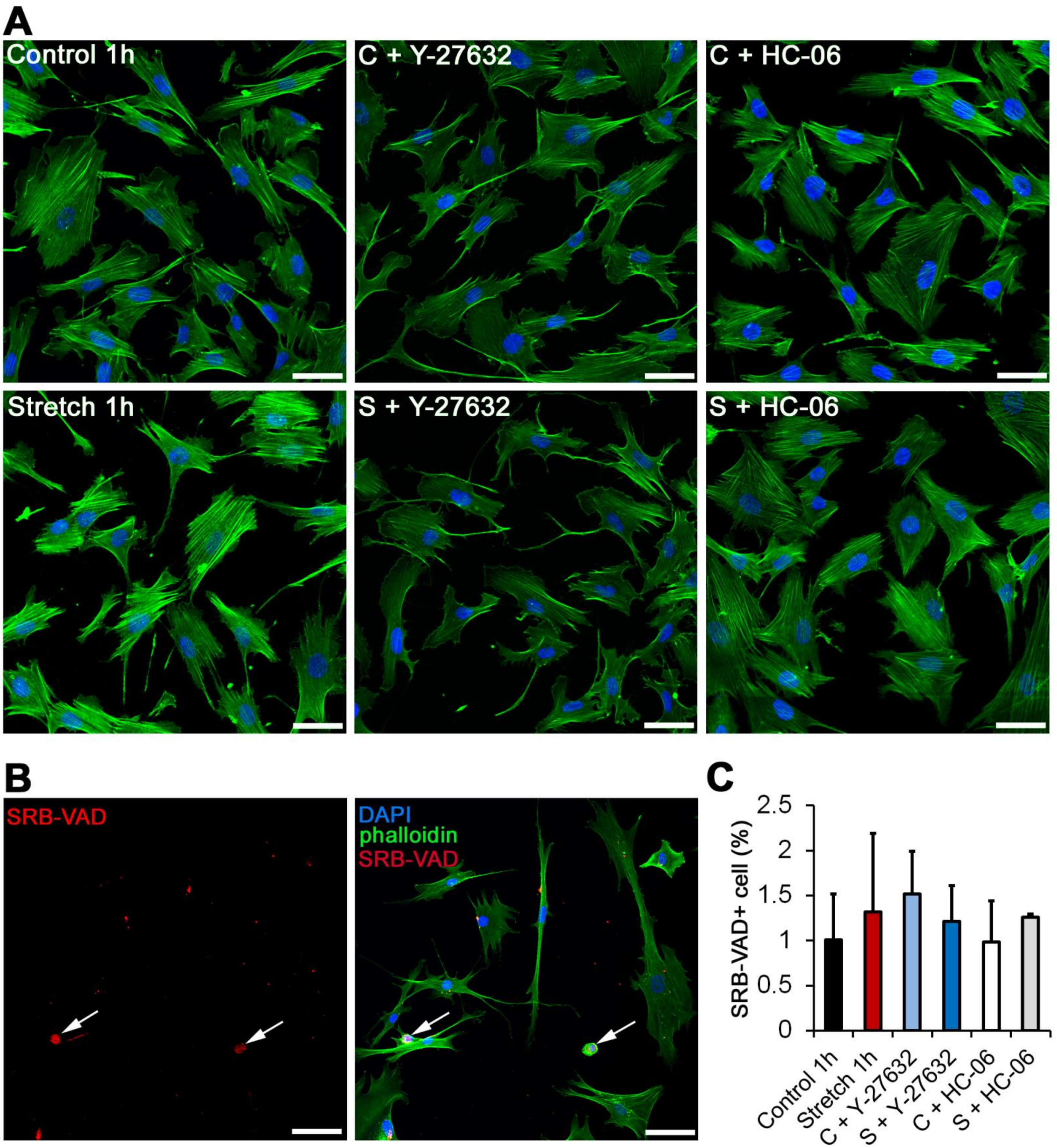

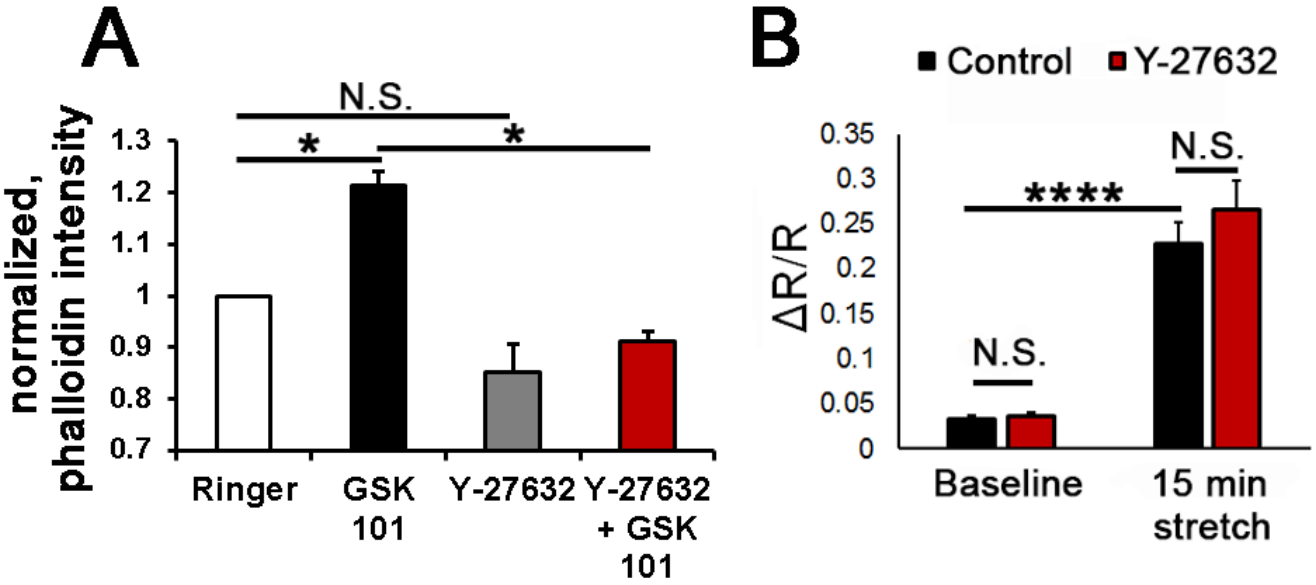

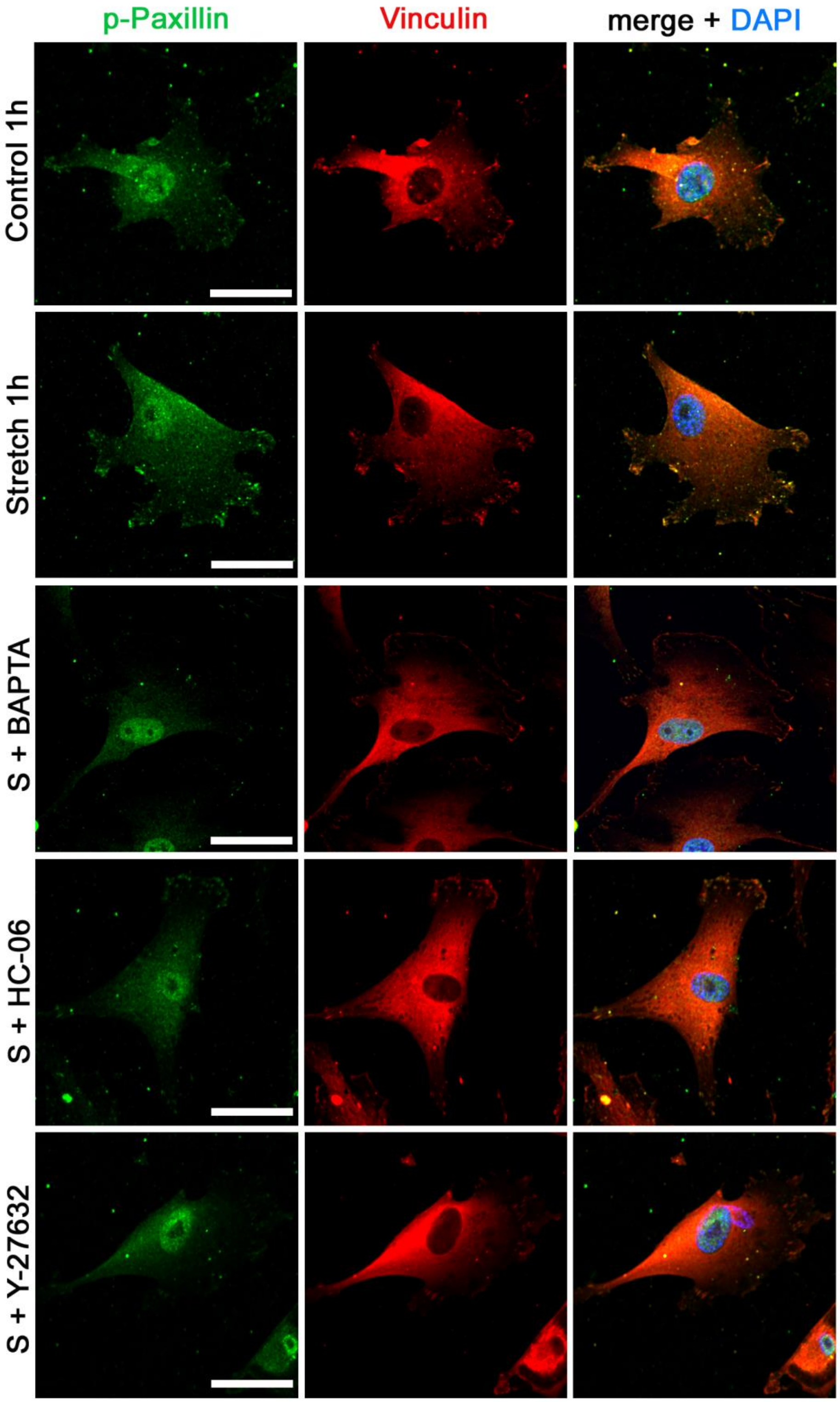

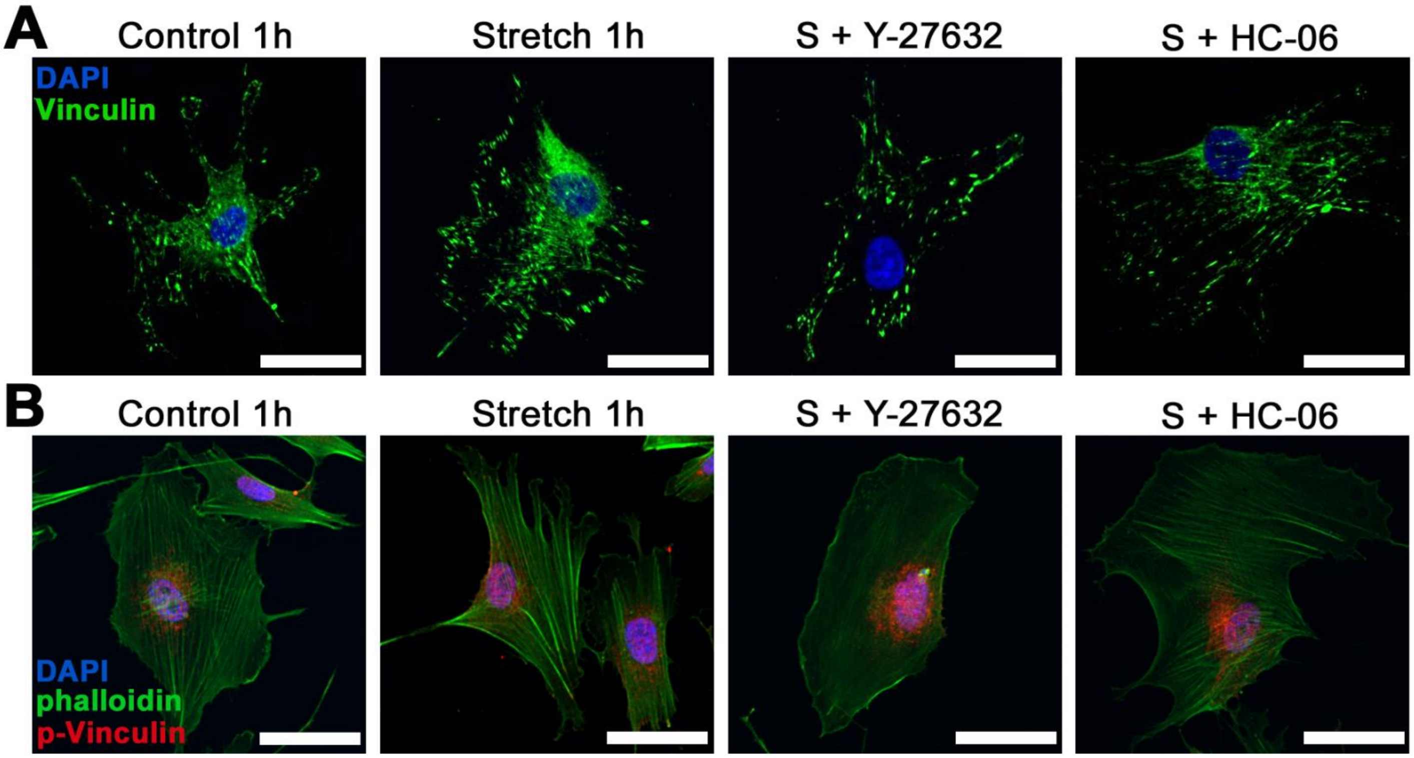

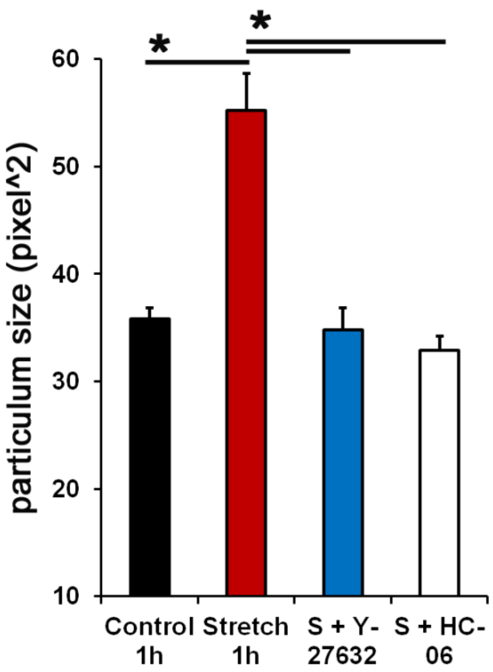

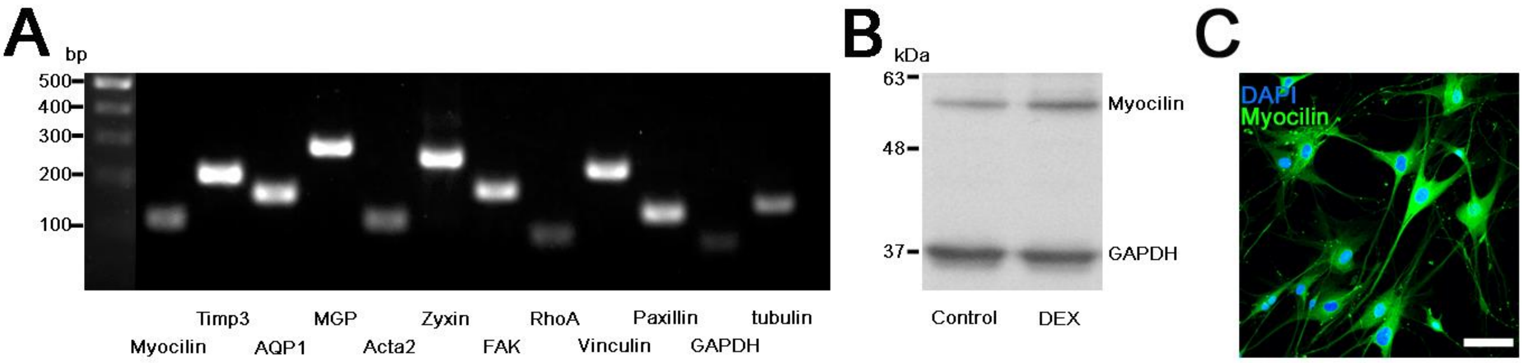

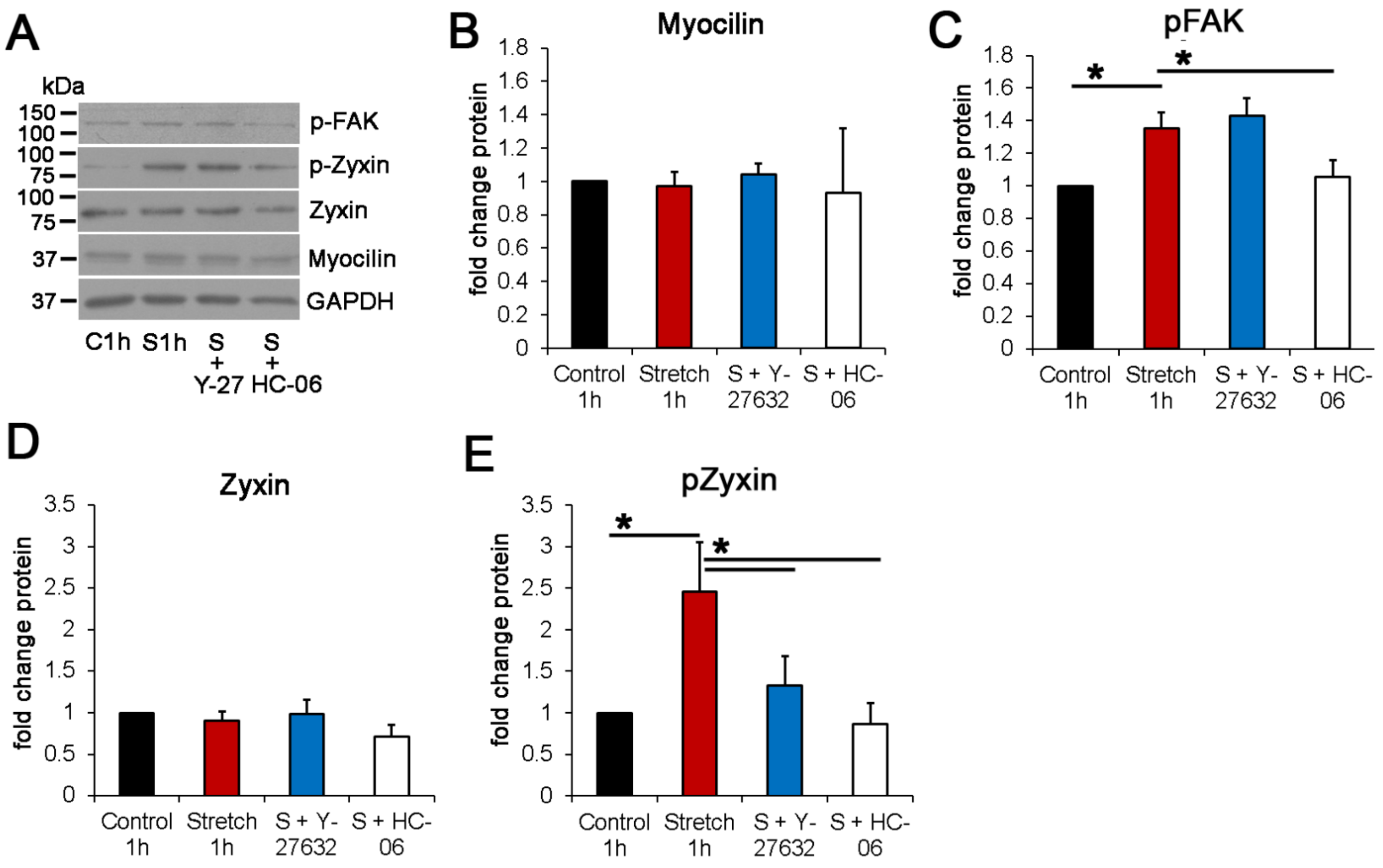
